# SiteCELL enables *on-site* PBMCs purification and cryopreservation for immune single cell profiling of diverse ancestries

**DOI:** 10.1101/2025.09.30.679317

**Authors:** Aarón Espinosa-Jaime, Octavio Zambada-Moreno, Jose Antonio Corona-Gomez, María de Jesús-Ortega, Marcela Hernandez-Coronado, Barbara Dos Santos Dias, Larissa Alvez Diniz, Laura Leaden, Guillherme Martelossi Cebinelli, Domenica Marchese, Adolfo Rojas-Hidalgo, Emiliano Vicencio, Diego Pérez-Stuardo, Sebastián Urquiza-Zurich, Marisol Espitia Fajardo, Alicia Colombo, Gerardo Donoso, Tamara Méndez, Carla Gallo, Hugo Guerrero-Cázares, Carla Daniela Robles-Espinoza, Patricia A. Possik, Guillermo Barreto, Lucía Spangenberg, Ricardo A. Verdugo, Vinicius Maracaja-Coutinho, Patricia Severino, Andrés Moreno-Estrada, Carlos Ortiz-Ramírez

## Abstract

Single cell genomics has improved our knowledge of immune function and heterogeneity. In recent years, the steady increase in the number of cells and individuals profiled as part of large multinational projects has enabled the characterization of cellular differences across human populations and ancestries. However, methods for collecting and processing peripheral blood mononuclear cells (PBMCs) for downstream single cell sequencing are difficult to implement in remote and rural settings. This has resulted in a lack of representation of underserved communities across the Global South in current initiatives. Hence, we developed SiteCELL, a method that enables purification and cryopreservation of PBMCs from whole blood at the site of collection using minimal laboratory equipment and without electricity. By comparing matched samples of purified PBMCs, we showed that SiteCELL performs as well as ficoll density gradient (FDG), both in laboratory and rural settings. This method ensures accurate recovery of cell type proportions and excels in reducing stress and minimizing variability across sampling batches. These advantages make it particularly well suited for implementation in challenging settings across countries, thereby enabling the inclusion of underrepresented ancestries in cellular atlases.

## Introduction

The advent of single cell technologies has enabled a deeper characterization of the immune system by disentangling its highly heterogeneous cellular and molecular responses in the context of health and disease. This higher resolution in immune profiling has already proven useful for the identification of new cellular subtypes and developmental checkpoints, as well as more precise biomarkers for disease states (*1-4*).

Although single cell studies were initially done by sourcing PBMCs from relatively small cohorts, several initiatives now exist that aim to build comprehensive cellular atlases including information from hundreds of individuals and millions of cells. These initiatives have opened up the possibility of correlating cell types and functions to biological conditions, as well as assessing the effect of demographics and ancestry on gene expression (*5, 6*). However, populations living in low to middle income or remote areas are largely absent from those projects due to limited access to specialized equipment and the trained personnel necessary to isolate and preserve PBMCs *on-site* (*7*). This is also true for many clinical settings in which samples cannot be processed promptly to avoid changes in gene expression.

In most studies, PBMCs are isolated from whole blood and then cryopreserved for future downstream analyses, including single cell profiling. However, the standard Ficoll Density Gradient (FDG) method for PBMC isolation consists of separating cells into distinct layers and recovering the PBMCs containing fraction (*8*). This can be time-consuming and requires specialized equipment not available in remote settings. Shipping fresh blood to a laboratory is not always an option because some sampling locations can be days away from the nearest city, and changes in gene expression have been detected in PBMCs after only four hours post blood draw (*9-11*). In addition, variability across sampling sites is common, especially regarding cell composition and the introduction of unwanted cells like platelets and erythrocytes.

Alternatives that avoid isolating PBMCs at the point of collection by either sample fixation or whole blood cryopreservation exist (*12-14*). However, fixation is not compatible with most single cell sequencing technologies, and isolating PBMCs from blood that has undergone a cycle of freezing and thawing is challenging and can introduce biases. For example, it has been reported that cell type proportions change after PBMCs isolation from thawed whole blood. In particular, fewer myeloid cells are recovered because they tend to attach to stressed or dying granulocytes, which are particularly sensitive to freezing and are targeted for removal during purification (*12*). A way to circumvent this issue was recently reported, in which the use of either FACS or enrichment of live PBMCs after initial purification using dead cell removal beads together with PE conjugated antibodies can render good quality cells for single cell downstream processing (*13*). However, this means extra processing steps and time, increased costs, and the need for FACS equipment, which is not available or affordable for many research institutions. In addition, isolation of PBMCs post-thawing means that they cannot be frozen again and stored in biobanks, a significant drawback for large-scale and cross-purpose studies.

We report an optimized method for *on-site* preservation of PBMCs, called SiteCELL, that allows simple and consistent PBMC purification and cryopreservation in remote or challenging settings. Using immunomagnetic negative selection and rate-controlled freezing in dry ice, PBMCs can be purified and cryopreserved at the site of collection in less than one hour for a single batch, without the need of specialized or electrically powered equipment. Purified PBMCs are cryopreserved *on-site* and transported in dry ice, which avoids any transcriptional perturbations or cell stress. The use of standardized checklists, the deliberate avoidance of electrical equipment, and the harmonization of reagent usage are key improvements that enhance the protocol’s robustness. Collectively, these optimizations facilitate the recovery of high-quality cells suitable for downstream single cell analysis. Furthermore, this method supports biobanking of PBMCs, which is particularly advantageous when clinical assays are conducted in parallel (*15*).

Here, we first tested SiteCELL under laboratory conditions and showed it performs equally well compared to the FDG gold standard in terms of cell viability, cell type relative proportion capture, and reduced gene expression changes. We then implemented SiteCELL in remote and rural settings to profile indigenous communities across Latin America, with the goal of increasing representation of Native Latin American ancestries in the generation of an immune single cell atlas by the LatinCells consortium (www.latincells.org). We show that SiteCELL maintains remarkable performance in challenging purification conditions, outperforming FDG in reduction of cell stress, cell purity (absence of platelets and erythrocytes), and reduced technical variability across batches, making it an ideal method for multi-national initiatives and for increasing representation of underserved communities.

## Results

### Analyses of matched samples demonstrate similar performance between FDG and SiteCELL

To evaluate our method’s performance, we first collected blood samples from healthy individuals at Hospital Israelita Albert Einstein (Sao Paulo, Brazil) and processed aliquots in parallel using both SiteCELL and FDG. In brief, samples were split into two and used to purify and cryopreserve PBMCs, followed by thawing and single cell profiling using the 10X Genomics Next GEM single cell 3’v3.1 HT kit. In the case of SiteCELL, negative selection of red blood cells, platelets and granulocytes was done with immunomagnetic beads, followed by addition of cryopreservation media (see detailed protocol) and controlled freezing (1 oC/min) in dry ice using a cool cell container (Fig. 1). For the rest of the blood sample, purification was done by gradient centrifugation, followed by the addition of cryopreservation media and controlled freezing in a -80°C ultrafreezer. Post-thaw, processing steps were the same for both methods and included multiplexing cells from four individuals in a single Chromium chip well. Genotype-based demultiplexing was applied to sequenced data, and analyses were performed to estimate data quality.

**Fig. 1.**
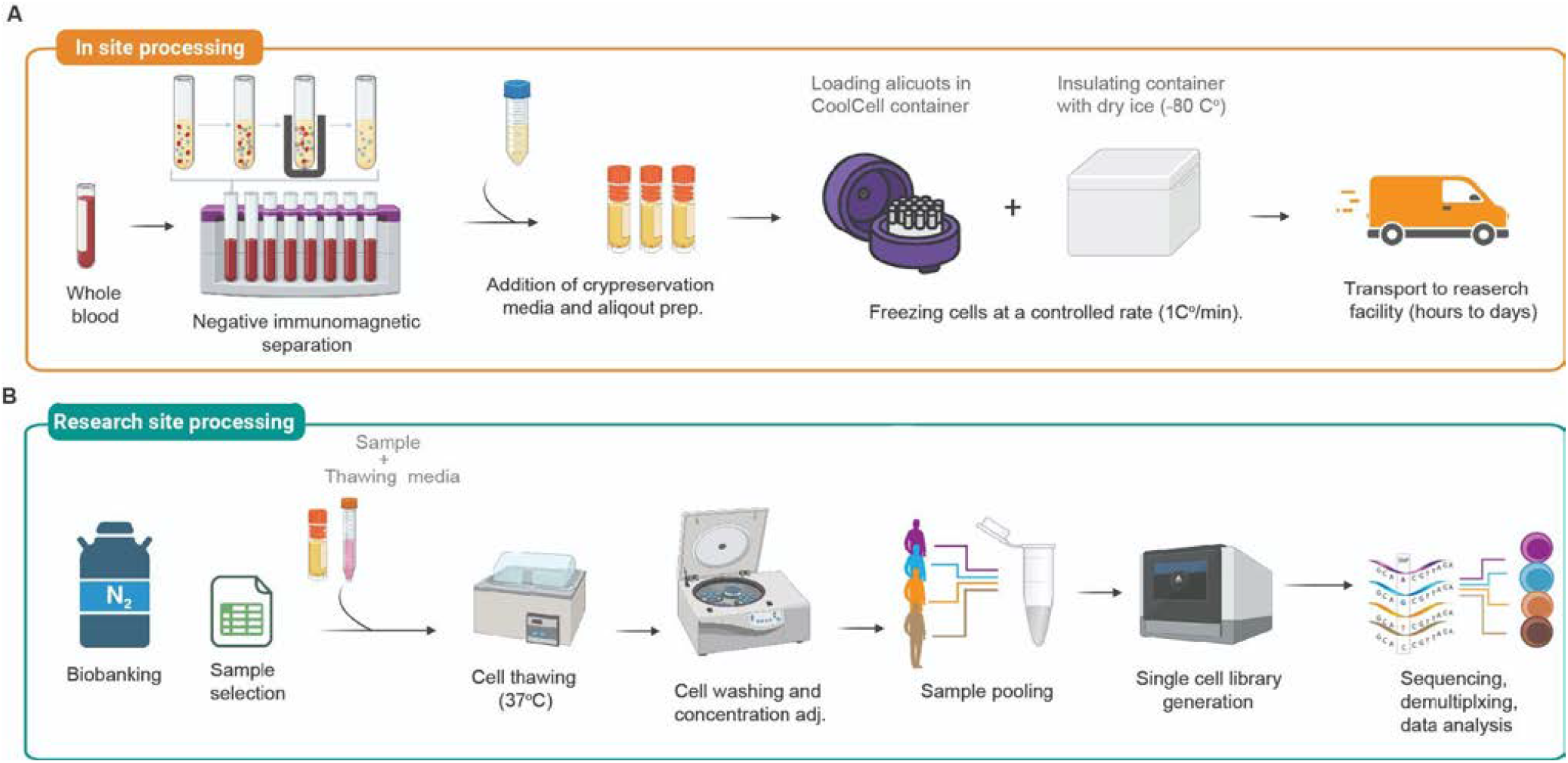
SiteCELL protocol workflow. **A)** Diagrams illustrating the individual steps performed at the site of collection to purify, cryopreserve, and transport frozen PBMCs to the nearest research facility. **B)** Workflow showing individual steps performed at the laboratory to thaw, wash, and prepare single cell libraries. With SiteCELL, PBMCs are transported frozen at -80 °C and readily stored in biobanks at the nearest research facility.

Cell viability upon thawing was approximately 90% for both methods, with slightly higher values observed for samples processed using SiteCELL. (93% ± 2.6 SiteCELL and 86% ± 3.1 FDG) (Fig. 2A). The difference between methods was statistically significant (p = 0.006, t-test).

**Fig. 2.**
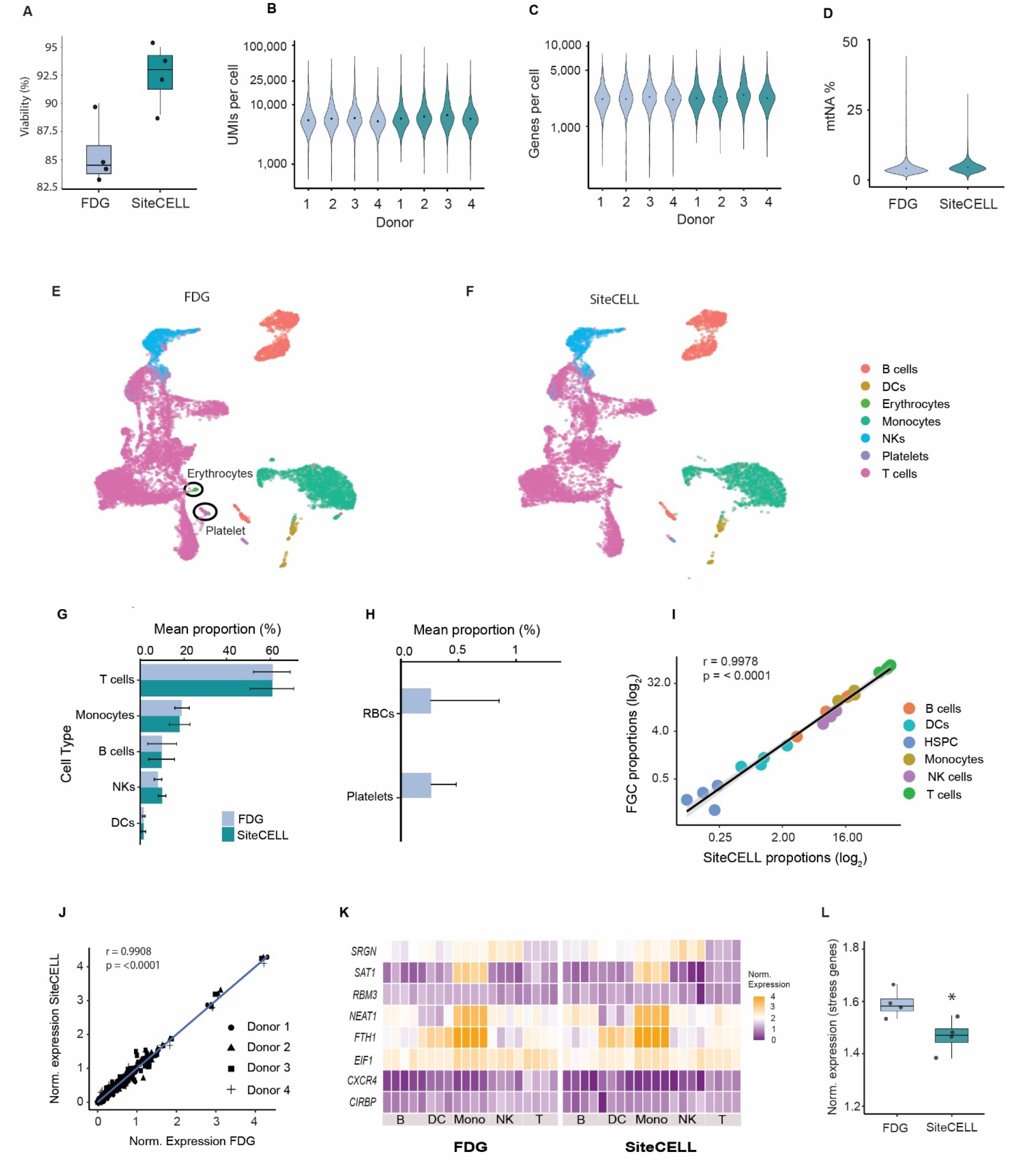
Comparison of matched PBMC samples processed with SiteCELL and FDG. **A)** Cell viability for samples processed by FDG (light grey) and SiteCELL (green) was quantified using trypan blue-stained dead cells. **B)** Number of UMIs per cell for each donor calculated by the Seurat package for FDG (light grey) and SiteCELL (green) samples. **C)** Number of genes per cell for each donor calculated by the Seurat package for FDG (light grey) and SiteCELL (green) samples. **D)** Percentage of mitochondrial transcripts from the total number of transcripts per cell across donors for FDG (light grey) and SiteCELL (grey) samples. **E)** UMAP showing cells colored by major immune cell types for FDG samples. The presence of platelets and erythrocytes is highlighted by dark circles. **F)** UMAP showing cells colored by major immune cell types for SiteCELL samples. **G)** Comparison of relative cell proportions of major immune cell types for FDG (light grey) and SiteCELL (green) samples. **H)** Comparison of the proportion of contaminating cell types, platelets, and erythrocytes per protocol. **I)** Correlation of cell type proportions between matched samples processed by FDG and SiteCELL. Each dot from each individual color represents a donor. **J)** Correlation of gene expression levels for cell identity genes between matched FDG and SiteCELL samples. Each dot represents a gene per donor. The blue line represents a linear regression fit. **K)** Heatmap showing normalized expression values for major immune cell types of eight stress-responsive genes. FDG samples are grouped at the left and SiteCELL samples at the right. **L)** Average values of the same eight stress-responsive genes per donor for FDG (light grey) and SiteCELL (grey) samples. Asterisk indicate statistical significant difference (T-test, p = 0.003)

In total, we sequenced 13,271 cells processed with gradient centrifugation and 10,620 processed with SiteCELL, each containing cells from the same four individuals. Both protocols produced high-quality data, with SiteCELL samples having slightly more Unique Molecular Identifiers (UMIs) (avg. 7056 ± 4598) and genes per cell (avg. 2484 ± 946) compared to FDG samples (UMIs avg. 6426 ± 3957, genes per cell avg. 2366 ± 868) (Fig. 2B & C). We estimated the percentage of mitochondrial RNA, as it is known to be high (>15%) in lysed cells or those undergoing apoptosis. Cells from both protocols showed a low average proportion of mitochondrial transcripts (≈ 4%) (Fig. 2D). However, FDG samples had a slightly higher number of cells with a mitochondrial transcript content above 15% (FDG 139 cells, SiteCELL 26 cells).

Biases in recovered cell type proportions have been reported for FDG alternative methods. Thus, to compare relative cell type proportions between matched samples, we clustered cells according to their transcriptome similarity using the Uniform Manifold Approximation and Projection (UMAP) analysis from the Seurat package. Cells from different individuals were integrated using Seurat’s CCA algorithm and annotated using reference-based mapping by Azimuth with I2 resolution. All cell types were identified in our PBMC samples with a similar distribution (Fig. S1). We calculated the mean relative proportion of major cell types (T cells, monocytes, natural killer cells, dendritic cells, and B cells) and found that they were in the expected range of what has been previously reported by single cell studies (*16*) (Fig. 2D, E, and F). Corroboration of this result was done by employing FACS analysis to quantify antibody-labeled cell types (Fig. S2). We quantitatively determined cell type proportion similarity between protocols by calculating Pearson correlation across parallel-processed samples. Correlation was very high (r = 0.997, p < 0.001), showing that proportions were almost identical and no cell capture bias was introduced by SiteCELL (Fig. 2I). Notably, we clearly detected the presence of platelets and red blood cells in samples processed by FDG but not in SiteCELL samples, indicating the latter clearly performs better in terms of purity (Fig. 2H).

It has been shown that moderate differences in sample handling and processing time can induce transcriptional changes even if cell viability seems unaffected (*9*). Hence, we analyzed the activity of the top marker genes for each cell subtype (a total of 219 genes) and performed correlation analyses between matched samples to determine if processing differences attributed to SiteCELL could result in expression biases. Correlation between protocols was very high (r = 0.99, p = 0.0001), showing that SiteCELL reliably conserves expression signatures of cell identity genes (Fig. 2J). Furthermore, we estimated cell stress due to processing by quantifying the expression levels of eight genes that are known to be upregulated in PBMCs in response to stress conditions and processing delays (*17*). Most genes showed slightly higher expression values across cell types in samples processed by FDG compared to SiteCELL, except for *NEAT1* (Fig. 2K). At the cell type level, natural killer cells displayed more differences between protocols, presenting more stress when subjected to FDG. We also noted that regardless of the protocol, monocytes showed the strongest evidence of stress and thus seem to be the most susceptible cell type to handling (Fig. 2K). Overall, when quantifying normalized expression of all stress-related genes across all cell types, SiteCELL processed cells showed significantly lower values (p = 0.003, t-test) (Fig. 2L). Since processing times were similar for both protocols under laboratory conditions, this result suggests that the extra centrifugation steps, combined with the exposure to Ficoll necessary for PBMCs isolation with FDG, might induce some stress.

### SiteCELL delivers accuracy in cell type proportions and optimal quality when implemented in remote settings

We showed that SiteCELL performs as well as FDG in the laboratory setting using matched samples. However, since the main goal of SiteCELL is to enable PBMC collection and purification in remote and rural locations, we next evaluated the performance and reproducibility of the protocol in the field. We analyzed three independent batches containing samples collected from six indigenous communities across Latin America, in which PBMCs were purified and cryopreserved *on-site*. Libraries were prepared by two independent processing hubs in México (two batches) and Chile (one batch). As a reference, we analyzed in parallel five publicly available PBMC datasets generated by different research teams using FDG and the same single cell library preparation technology (10x Genomics) (*18-22*). We assessed sample quality focusing on three key criteria: transcript capture, relative cell proportion accuracy, and cell stress. Ancestry and sampling details are presented in Tables 1 and 2.

**Table 1.** LatinCells sample IDs (SC = SiteCELL).

| Dataset | Donor label | Ancestry | Location | Country | Viability (%) |
| --- | --- | --- | --- | --- | --- |
| SC batch 1 | LCMX-0142 | Otomí | Guanajuato | Mexico | 94.7 |
| SC batch 1 | LCMX-0143 | Otomí | Guanajuato | Mexico | 92.45 |
| SC batch 1 | LCMX-0144 | Otomí | Guanajuato | Mexico | 90.2 |
| SC batch 2 | LCMX-0004 | Maya | Yucatán | Mexico | 97.9 |
| SC batch 2 | LCMX-0147 | Otomí | Guanajuato | Mexico | 99.1 |
| SC batch 2 | LCCO-0003 | Piapoco | Vichada | Colombia | 97.9 |
| SC batch 2 | LCCO-0025 | Pastos | Nariño | Colombia | 79 |
| SC batch 3 | LCCL-0089 | Aymara | Iquique | Chile | 93 |
| SC batch 3 | LCCL-0091 | Aymara | Iquique | Chile | 93 |
| SC batch 3 | LCUY-0001 | Admixed | Montevideo | Uruguay | 95 |
| SC batch 3 | LCUY-0002 | Admixed | Montevideo | Uruguay | 93 |

**Table 2.**
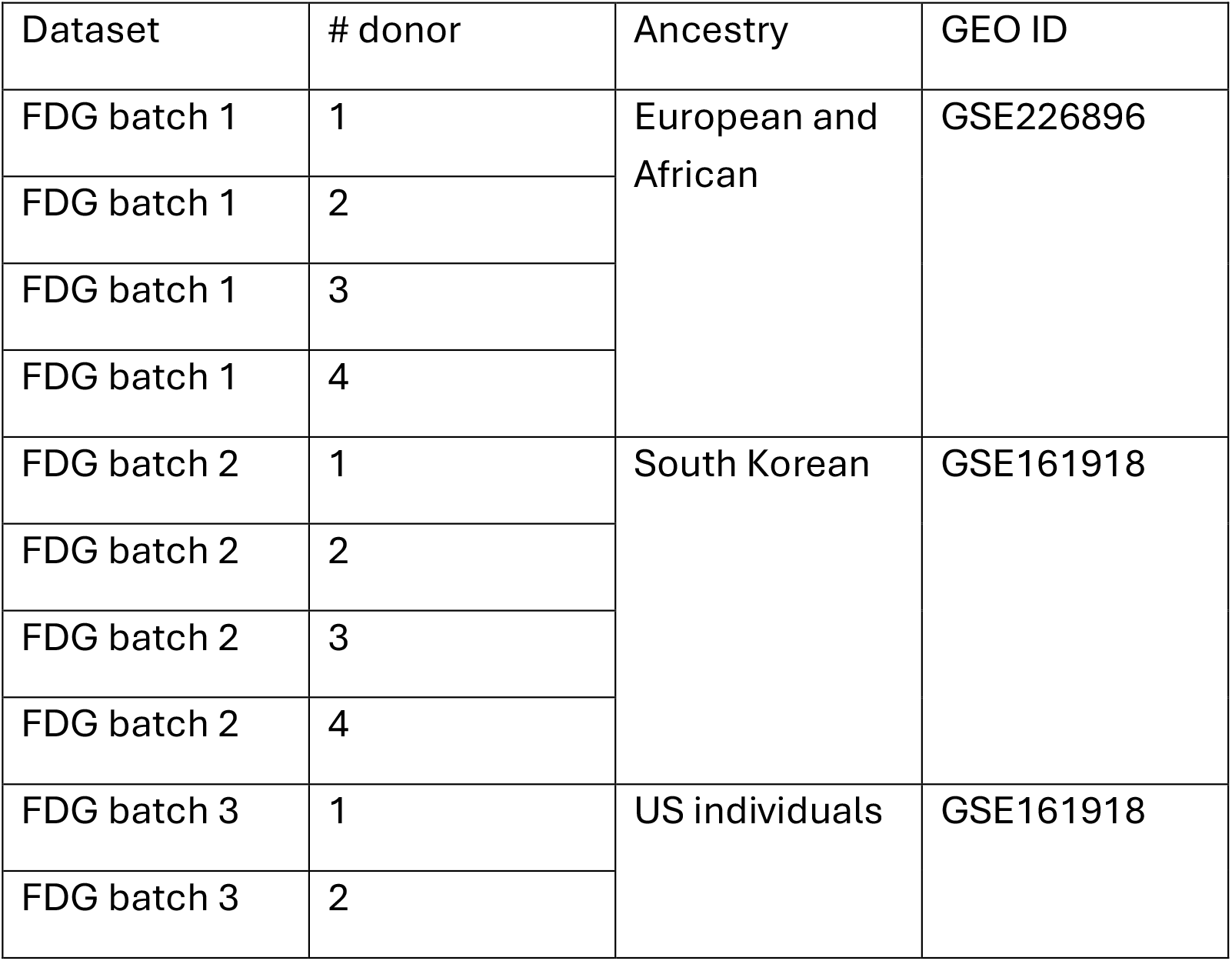
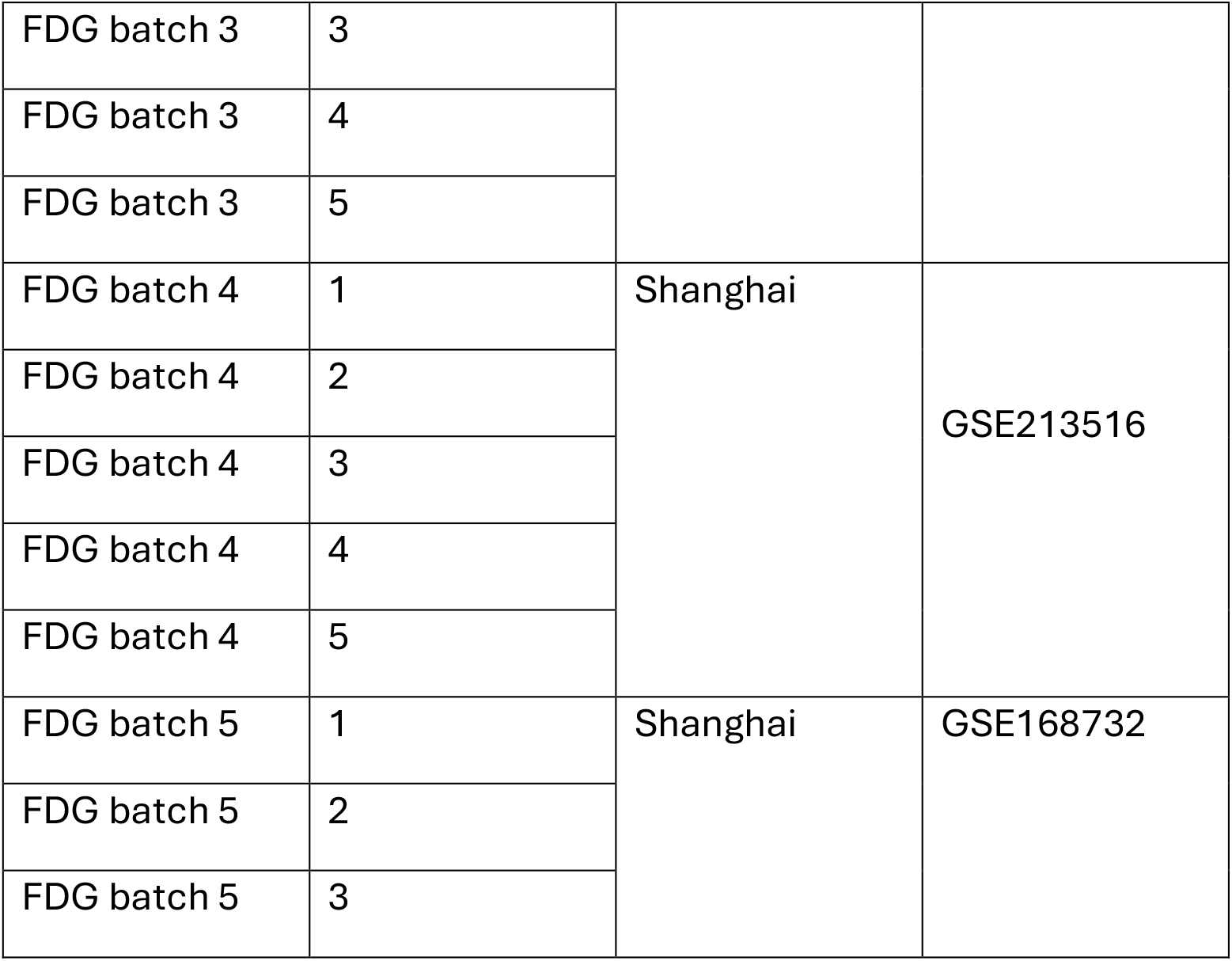
Ficoll datasets and GEO IDs.

For the SiteCELL method, post-thaw cell viability was above 90% for most samples, as reported before, except for a single Colombian sample (Table 1). A total of 63,992 cells were sequenced in batches of 2-4 individuals. Analyses showed samples had a high average of genes (2,664 ± 340) and unique UMIs per cell (7,796 ± 1487). In the case of FDG datasets, viability was not reported for most batches. Each dataset had cells from 3-5 individuals, and a total of 220,604 cells were analyzed. A significantly lower average of genes (1,388 ± 260) and unique UMIs (4,424 ± 1,143) per cell were detected compared with SiteCELL (Fig. 3A & B). This was not due to differences in sequencing depth, because FDG samples had a higher number of reads per cell (Table 3). However, we note that although both SiteCELL and FDG libraries were prepared using 10X Genomics single cell RNAseq technology, some of the latter were processed using older versions of that kit, which might have reduced transcript capture efficiency. Detailed information for both datasets can be found in Table 3.

**Table 3.** Datasets metadata. Includes a brief description of the number of cells, number of genes and number of UMIs (SC = SiteCELL).

| Dataset | Processing Hub | # donor | Donor label | Total Cells | Total cells in dataset | average UMI | dataset average UMI | average genes per cell | dataset average genes per cell | dataset sequencing depth |
| --- | --- | --- | --- | --- | --- | --- | --- | --- | --- | --- |
| SC batch 1 | Mexico | 1 | LCMX-0142 | 5209 | 34989 | 9748 | 9842 | 3138 | 3147 | 70,000 reads per cell |
| SC batch 1 | Mexico | 2 | LCMX-0143 | 8402 |  | 9232 |  | 3066 |  |  |
| SC batch 1 | Mexico | 3 | LCMX-0144 | 19957 |  | 10104 |  | 3181 |  |  |
| SC batch 2 | Mexico | 1 | LCMX-0004 | 3992 | 16664 | 6674 | 6381 | 2394 | 2368 | 36,000 reads per cell |
| SC batch 2 | Mexico | 2 | LCMX-0147 | 6162 |  | 6756 |  | 2468 |  |  |
| SC batch 2 | Mexico | 3 | LCCO-0003 | 2567 |  | 5677 |  | 2206 |  |  |
| SC batch 2 | Mexico | 4 | LCCO-0025 | 3332 |  | 5898 |  | 2283 |  |  |
| SC batch 3 | Chile | 1 | LCCL-0089 | 2714 | 12339 | 7990 | 7931 | 2737 | 2654 | 32,000 reads per cell |
| SC batch 3 | Chile | 2 | LCCL-0091 | 2379 |  | 8194 |  | 2733 |  |  |
| SC batch 3 | Chile | 3 | LCUY-0001 | 2655 |  | 7372 |  | 2515 |  |  |
| SC batch 3 | Chile | 4 | LCUY-0002 | 3030 |  | 8120 |  | 2584 |  |  |
| FDG batch 1 | Europe | 1 | 5 | 5901 | 23142 | 5302 | 3681 | 1598 | 1340 | 300,000 reads per cell<br>?*** |
| FDG batch 1 |  | 2 | 14 | 5780 |  | 6490 |  | 1798 |  |  |
| FDG batch 1 |  | 3 | 13 | 6196 |  | 5149 |  | 1522 |  |  |
| FDG batch 1 |  | 4 | 19 | 5265 |  | 7215 |  | 2000 |  |  |
| FDG batch 2 | South Korea | 1 | Immuno1 | 14785 | 60982 | 3587 | 5993 | 1408 | 1719 | 50,000 reads per cell |
| FDG batch 2 |  | 2 | Immuno2 | 22792 |  | 2796 |  | 1095 |  |  |
| FDG batch 2 |  | 3 | Immuno3 | 13216 |  | 3749 |  | 1392 |  |  |
| FDG batch 2 |  | 4 | Immuno4 | 10189 |  | 5711 |  | 1719 |  |  |
| FDG batch 3 | US | 1 | CHI014 | 20114 | 77501 | 3498 | 3517 | 1090 | 1094 | NA |
| FDG batch 3 |  | 2 | SHD1 | 13923 |  | 3513 |  | 1095 |  |  |
| FDG batch 3 |  | 3 | SHD3 | 16006 |  | 3502 |  | 1094 |  |  |
| FDG batch 3 |  | 4 | SHD5 | 15379 |  | 3521 |  | 1091 |  |  |
| FDG batch 3 |  | 5 | SHD6 | 12486 |  | 3563,7 |  | 1106,4 |  |  |
| FDG batch 4 | Shanghai | 1 | HC1 | 8019 | 42771 | 4716 | 4850 | 1280 | 1405 | 110,000 reads per cell |
| FDG batch 4 |  | 2 | HC2 | 8279 |  | 4931 |  | 1395 |  |  |
| FDG batch 4 |  | 3 | HC3 | 9848 |  | 4520 |  | 1547 |  |  |
| FDG batch 4 |  | 4 | HC4 | 7581 |  | 5636 |  | 1612 |  |  |
| FDG batch 4 |  | 5 | HC5 | 9944 |  | 4116 |  | 1228 |  |  |
| FDG batch 5 | Shanghai | 1 | kawasaki 1 | 5690 | 16208 | 4110 | 3876 | 1388 | 1359 | 69,000 reads per cell |
| FDG batch 5 |  | 2 | kawasaki 2 | 5077 |  | 3625 |  | 1288 |  |  |
| FDG batch<br>5 |  | 3 | kawasaki<br>3 | 5441 |  | 3667 |  | 1396 |  |  |

**Fig. 3.**
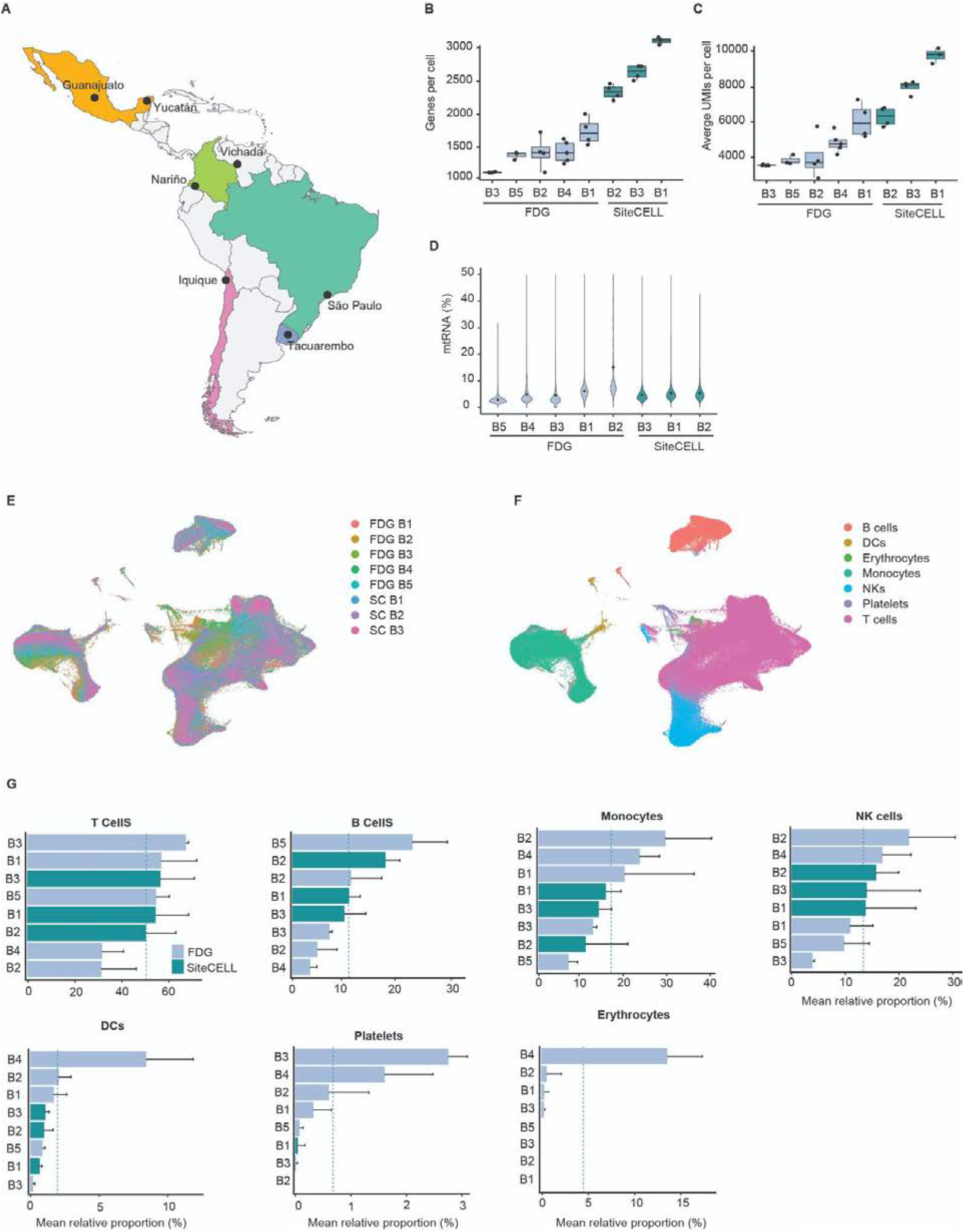
SiteCELL performance for PBMCs isolation in remote and rural settings. **A)** Average values of genes per cell for individual samples and across batches calculated with the Seurat package for FDG (light grey) and SiteCELL (green). **B)** Average values of UMIs per cell for individual samples and across batches calculated with the Seurat package for FDG (light grey) and SiteCELL (green). **C)** Percentage of mitochondrial transcripts from the total number of transcripts per cell across batches for FDG (light grey) and SiteCELL (green) samples. **D)** UMAP showing cells colored by sample processing batches. FDG = ficoll density gradient, SC = SiteCELL. **E)** UMAP showing cells colored by major immune cell types for all integrated batches with the CCA algorithm from the Seurat package. **F)** Comparison of relative cell proportions of major immune cell types for FDG (light grey) and SiteCELL (green) samples across batches.

The percentage of mitochondrial transcripts per cell was evaluated for all datasets. Most had an average below 10%, which is considered acceptable, except for the FDG batch 2. Overall, there was a higher variability across the FDG datasets, with mitochondrial transcripts averages ranging from 2.9% to 16% (Fig. 3C). Values for SiteCELL datasets ranged from 5.0 to 5.43%. We also calculated the number of cells with more than 15% of mitochondrial RNAs, which are considered to be severely stressed or dying, and observed a substantial difference between protocols. For SiteCELL, we found that 1.7% of all cells were above this threshold (1,105 total cells), while for FDG datasets, 24% of all cells had more than 15% of mitochondrial transcripts (11,219 total cells). Again, there was substantial variability across datasets, with FDG batches 1 and 2 contributing most cells with a high proportion of mitochondrial transcripts (> 15%).

To evaluate whether our protocol introduced any bias in cell type capture efficiency when applied to challenging settings (fieldwork conditions), we identified cell type populations in all datasets as described before. We clustered cells and integrated different batches using UMAP and CCA algorithms from Seurat, followed by annotation with Azimuth (Fig. 3D & E). Subsequently, cell type relative proportions were calculated for each major group across all datasets and protocols. Relative proportions for both SiteCELL and FDG samples were similar and matched previously reported values (*16*), with T cells comprising 65-70%, B cells 5-22%, NK 5-14% and monocytes 4-12% of the total number of cells (Fig. 3F). We also confirmed that SiteCELL samples had almost no platelet (5 platelet total counts) and no red blood cell contamination compared with FDG samples (Fig. 3F).

We further assessed if SiteCELL was able to accurately capture and report rare or low-abundant immune cells. We focused on T cells because they group some of the rarest PBMC subtypes, like CD8 TCM and CD4 CTL. For performing comparative analyses, T cells from both protocols were re-clustered to annotate subtypes, and their proportions were calculated relative to the total number of T cells (major group) (Fig. 4A & B). SiteCELL samples had the same number of T cell subtypes and in equal proportions (no statistical differences) compared to FDG samples, including the ones with a lower abundance: proliferating T, CD4 CTL, Treg, MAIT, CD8 TCM, gdT, and dnT (Fig.4C). The subtype that showed the biggest difference between protocols, in terms of total cell number, was CD8 TCM. However, most of the cells reported for FDG were contained in only one batch; hence, this was not statistically significant. Overall, no bias was found in rare subtype recovery when SiteCELL was applied.

**Fig. 4.**
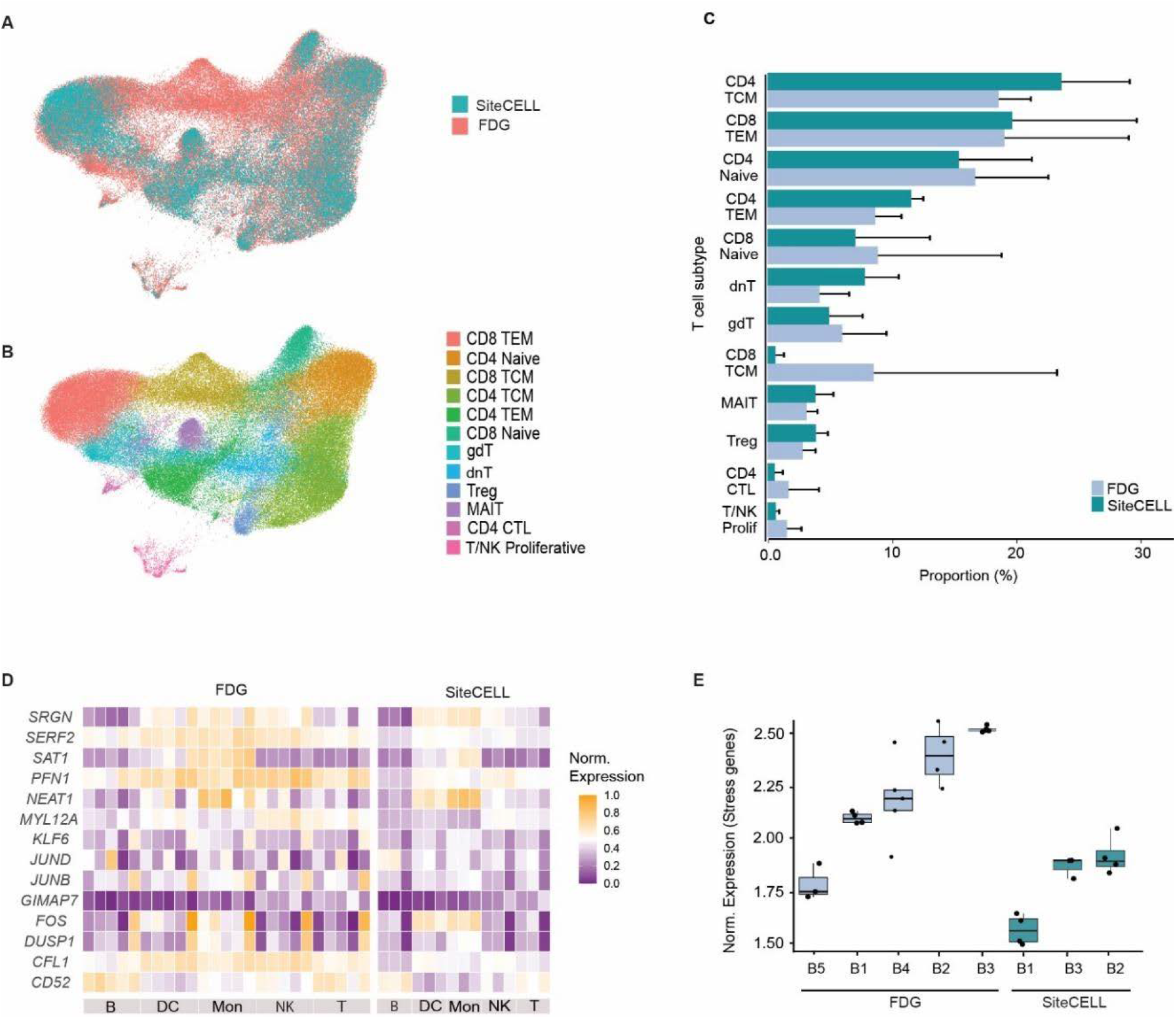
Recovery of low-abundant immune cell types and induction of stress genes. **A)** UMAP showing re-clustering of T cells of both FDG (light grey) and SiteCELL (green) samples. Cells are colored by protocol. **B)** UMAP shows re-clustered T cells from both protocols, in which cells are colored by identity subtypes. **C)** Comparison of relative T cell subtype proportions for FDG (light grey) and SiteCELL (green) samples across batches. **D)** Heatmap showing normalized expression values for major immune cell types of 12 stress-responsive genes. Samples processed by FDG are grouped at the left and SiteCELL samples at the right. **E)** Average values of the same 12 stress-responsive genes across batches processed by FDG (light grey) and SiteCELL (grey).

Regarding cell stress, we quantified the normalized expression of established stress-responsive genes as done previously. In this case, PBMC purification and cryopreservation were done in rural settings where sampling can be unpredictable, hence we also included genes from the JUN and FOS families and from the DUSP and GIMAP families. These are known to be upregulated when there are considerable delays before processing (*23*). SiteCELL samples showed low levels of stress-responsive gene expression compared to the FDG references (Fig.4D & E). Again, a cell type-dependent response was observed, with monocytes being the most affected regardless of the protocol, and NK cells showing more differences between protocols: very low or no expression in SiteCELL samples, to moderate expression in FDG samples. Overall, these findings highlight the suitability of SiteCELL to process samples in remote locations without inducing significant cell stress.

### SiteCELL improves consistency and robustness across sampling sites

Large-scale single cell studies with hundreds of samples collected across geographically diverse settings are becoming more common due to their broad applications. These projects are usually developed by consortia of multinational teams that collect and process samples across countries at different time points. Thus, ensuring data reproducibility is a major challenge. In this context, a method that is highly robust and reproducible is necessary to reduce batch effects across sites.

We evaluated variability across sample batches collected and processed with SiteCELL at different locations and time points, focusing on changes in cell type proportions and gene expression, and compared them to the FDG reference. For these analyses, we calculated the relative proportions of 27 cell subtypes and evaluated the consistency in which these were recovered. Variability was observed at the level of individuals, which was expected since it has been reported that age, sex, and overall health can produce moderate differences in immune cell type proportions (*24, 25*) (Fig. 5A). However, when assessing variability across batches, differences were apparent between protocols. To quantify this, we calculated the coefficient of variation from the average proportion of each cell subtype across batches. We observed that although the degree of variability is cell-type specific for both protocols, those cells processed with SiteCELL have an overall lower relative variability (≈ 40%) compared to those purified with FDG (≈ 95%), with one exception: B intermediate cells (Fig. 5B). This suggests that implementation of SiteCELL reduces batch effects due to processing. Notably, some cell types had a relatively high variability regardless of the protocol (CD8 naïve, CD4 naïve, and CD4 CTL) (Fig. 5B). Therefore, caution is advised when making conclusions about proportion differences of these subtypes due to ancestry or biological conditions. Altogether, these results support that SiteCELL preserves cellular composition even in challenging processing settings.

**Fig. 5.**
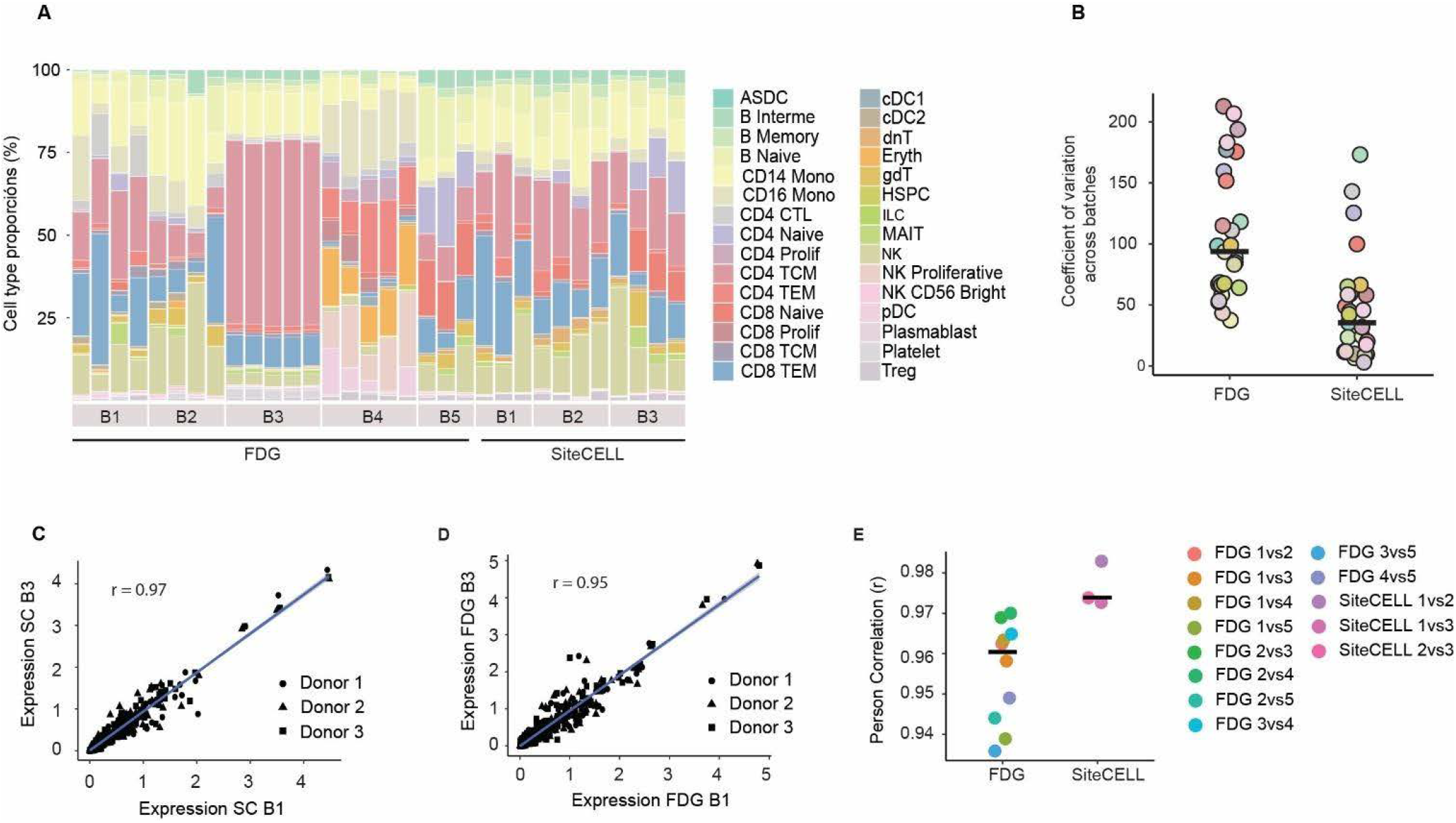
Cell subtype proportion and gene expression variability across sampling batches. **A)** Stacked bar plot showing relative cell proportions colored by immune subtypes across batches. Each stacked bar corresponds to a single individual. **B)** Coefficient of variation representing the relative dispersion of data (cell type proportion) for individual cell subtypes across batches. Higher values indicate higher variability across processing batches. Cell subtypes are colored as in A. **C)** Correlation of gene expression levels for cell identity genes between representative batches processed by SiteCELL (left) and FDG (right). Each dot represents a gene per donor. The blue lines represent linear regression fits. **E)** Pearson correlation values for all pair-wise comparisons of available batches for each protocol.

Finally, we also assessed variability in the expression levels of 219 cell identity genes (top markers for each cell subtype) across batches. We did pair-wise linear regression analyses and calculated Pearson correlation between all batches in each protocol. In general, gene expression correlation was high for both methods, suggesting that expression levels of cell identity genes are maintained even if batches are processed by different teams and at different time points. Although SiteCELL batches were collected and processed in remote and challenging settings, they had slightly higher correlation values (Fig. 5C & D) compared to FDG samples. SiteCELL *r* values across dataset comparisons ranged from 0.972 to 0.985, while FDG *r* values across datasets ranged from 0.936 to 0.97 (p = 0.001). Average correlation was *r* = 0.973 for SiteCELL samples and *r* = 0.961 for FDG (Fig. 5E). Hence, we conclude that SiteCELL is a robust and easy-to-implement method for PBMCs purification that reduces technical sample variability across sites. This makes it ideal for large-scale multinational single cell genomics projects and enables the inclusion of underrepresented human populations that live in remote and rural areas with no infrastructure.

## Discussion

Initiatives to build comprehensive immune cellular atlases are currently underway. However, most methods to purify and cryopreserve PBMCs are difficult to implement in remote locations that lack basic infrastructure, resulting in a lack of representation of underserved communities and ancestries. So far, the alternative to FDG for sampling in challenging settings has been the cryopreservation of whole blood samples immediately after blood draw without purification (*12, 13*). However, there are some drawbacks to freezing whole blood first and purifying PBMCs later. Mainly biases in cell type proportions, incompatibility with biobank storage, increased isolation complexity and costs, and induction of cell stress. The latter is especially relevant in the case of granulocytes, which become much harder to remove post-thawing due to loss of viability. To solve these issues, we developed SiteCELL which enables the purification and cryopreservation of PBMCs at the point of collection without electricity and minimal equipment. We employ a pre-freeze PBMC purification strategy that avoids cell stress and complicated granulocyte removal procedures after thawing. Cryopreserved PBMCs are transported frozen to the laboratory settings, where they can be immediately used for single cell profiling or directly stored in biobanks.

We analyzed matched samples processed to generate single cell data and demonstrated that SiteCELL performs as well as the gold standard FDG method in terms of cell quality and viability. Additionally, SiteCELL faithfully maintains PBMC cell type relative proportions as shown by the very high correlation scores of matched samples. This has been a drawback for some whole blood cryopreservation methods that report biases in relative cell type proportions due mainly to the fact that monocytes tend to attach to platelets or stressed granulocytes, which are targeted for removal post-thaw (*12*). Granulocytes tend to present low viability in whole blood freezing protocols because they are very sensitive to freezing and thawing cycles; hence, their removal at the point of collection before freezing is a considerable advantage of the SiteCELL protocol (*23, 26*). We also showed that SiteCELL is better at removing platelets and red blood cells, which is important because platelets can act as suppressors, inhibiting proliferative immune cells and affecting gene expression (*27, 28*). In addition, their unwanted presence implies a waste of valuable resources, since they can end up taking expensive single cell reagents and, in some cases, a considerable amount of sequencing reads.

In terms of gene expression, both methods show a very high correlation in the expression level of 219 cell subtype marker genes, showing that SiteCELL faithfully conserves expression profiles of genes related to cell identity. An important finding was that SiteCELL samples showed reduced levels of stress-responsive gene activation. This was especially evident for NK cells. Reduction was more significant when comparing multiple independent batches than in matched experiments. This could be due to a faster processing time with SiteCELL compared to other protocols. For example, cells can be immediately processed at the site of blood draw without the need to transfer them to any facility, and a batch can be purified and cryopreserved in less than an hour. This considerably reduces the time cells are exposed to *ex vivo* conditions, especially when sampling is conducted far from the laboratory. Another explanation could be that SiteCELL does not rely on centrifugation steps or ficoll plaque reagent, which can induce cell stress and toxicity when the gradient is not formed optimally. Absence of such processing steps might decrease the chances of cells being exposed to harsh conditions.

As large-scale single cell projects profiling hundreds or thousands of individuals and samples across contrasting settings become more common, simple but robust purification and cryopreservation methods are highly needed. We demonstrate that data generated from independent batches processed by SiteCELL at different rural locations was highly consistent. Two aspects were analyzed: cell type relative proportions and gene expression profiles of marker genes. In both cases, less variability was quantified across SiteCELL batches than across batches generated using FDG. However, we note that PMBC isolation with SiteCELL was highly harmonized across sampling sites, while FDG datasets used for comparison were generated by independent research groups which might have used slightly different versions of the FDG protocol. In any case, SiteCELL has a similar performance compared to FDG in terms of consistency across sample batches, even when deployed in rural and remote settings. We believe this is due to the relatively easy implementation of the immunomagnetic-based purification procedure employed by SiteCELL. In comparison, FDG can be more time-consuming and laborious as the process of layering and recovering the sample requires significant skill and can be difficult to master. Hence, another advantage of SiteCELL is that less training is required to achieve good purification performance. Overall, results showed this method is well-suited for deployment in remote and rural settings, facilitating inclusion of underrepresented populations in biomedical research by enabling global, multi-ethnic, single cell transcriptomic studies.

## Supporting information

SiteCELL bench-ready manual

## Acknowledgements

We thank all participants in this study as well as the community leaders for their support across all sampling sites. We acknowledge the Chang Zuckerberg Initiative (CZI) grant 2021-240108 (5022) and to the Secretaria de Ciencia, Humanidades, Tecnología e Inovación (CECIHTI) for postgraduate grants.

## Author contributions

A.E.J., O.Z.M. and C.O.R collected and processed PBMC samples, analyzed the data and helped write the manuscript. J.C.G., M.J.E., M.H.C, B.S.D., L.A.D., L.L., G.M.C., D.M., A.R.H., M.E.F., E.V., D.P.S., A.C., G.D, T.M. and S.U.Z. collected and processed PBMC samples. C.G., H.G.C., C.D.R.E., P.P., G.B., L.S., R.A.V., V.M.C.M, P.S, and A.M.E. were involved in community engagement, sampling logistics and manuscript edits. AM.E. and C.O.R. conceived experiments and helped write the manuscript.

## Supplemental figures

**Figure S1.**
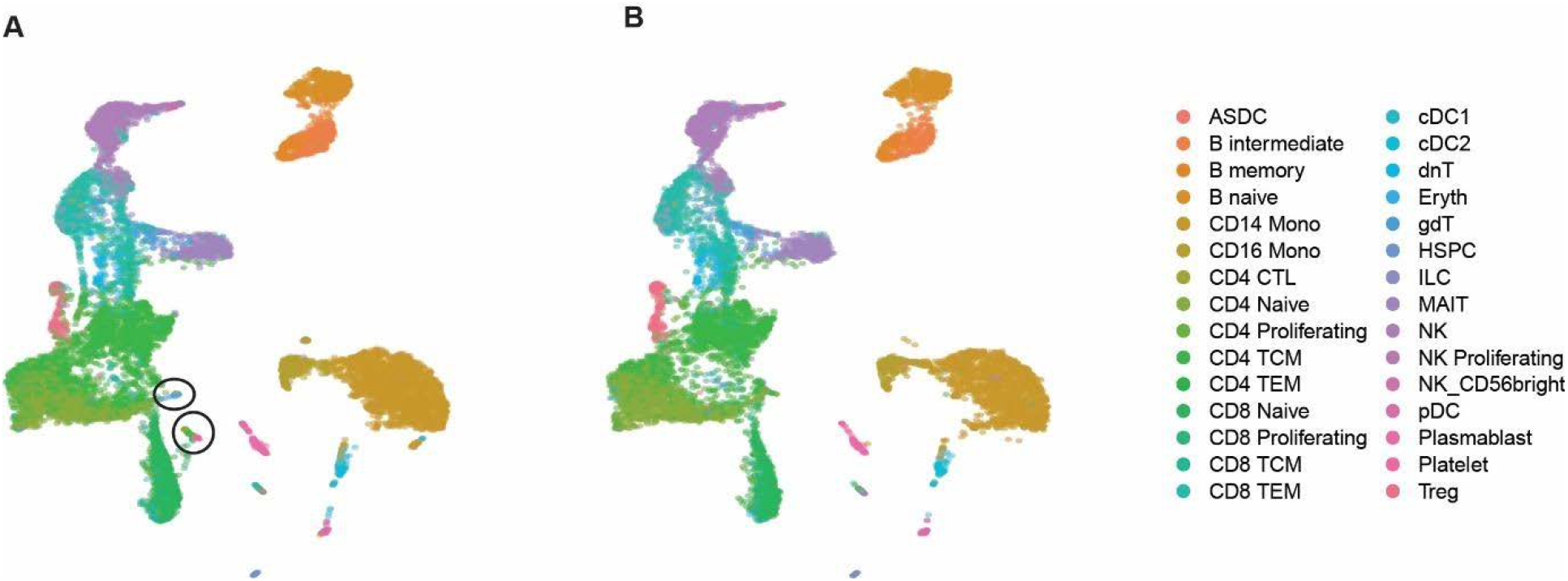
Immune cell subtype identification of matched samples processed by FDG and SiteCELL. **A)** UMAP shows cells colored by identity type for samples processed by FDG. 30 cell subtypes were identified, including platelets and erythrocytes. **B)** UMAP shows cells colored by identity type for samples processed by SiteCELL. 28 cell subtypes were identified, while platelets and erythrocytes were not found in these samples.

**Figure S2.**
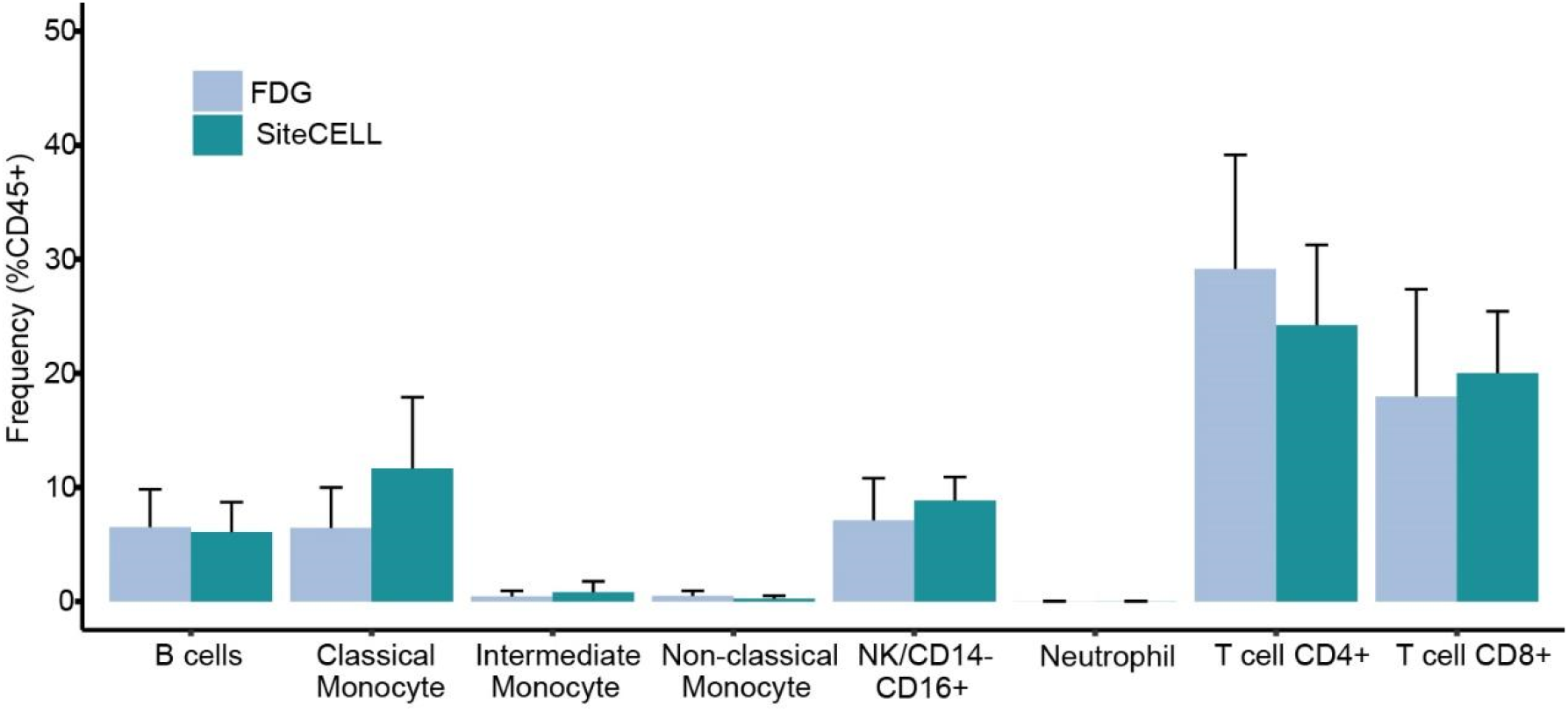
Cell type proportions determined by antibody cell labeling and FACS quantification. Matched samples were processed in parallel by FDG and SiteCELL, and cell type proportions qualified cell type-specific fluorescent-labeled antibodies and FACS. No statistical differences were found between protocols.

## Supplemental files

*On-site* PBMCs processing protocol V4.pfd

## Methods

### Participants Selection

Participants were selected based on the following inclusion criteria: they were adults, had not experienced an infectious process within the two weeks before participation, and had no prior diagnosis of autoimmune diseases. In some cases, self-reported indigenous ancestry was required. Written informed consent was obtained before participation in the study under protocol approved by each country (Brazil: CAAE 68371323.3.0000.0071, Chile: CEISH/091-2023/041, Colombia: CI 085-022, México: COBISH/085/2022, Uruguay: CEI 006-2023).

### Data Collection

For all participants included in the project, the following data was collected:

1. **Informed Consent:** A signed document confirming the participant’s voluntary and informed participation in the study.
2. **Questionnaire:** A comprehensive document used to gather general information about the participant. This included: personal details (full name, age, and contact information), family medical history, biometric data (weight, height, and blood type), prevalence of diseases, ancestry information.

These data were securely stored and used solely for this research, following ethical guidelines and ensuring participant confidentiality.

### Sample collection

Two types of biological samples were collected from each participant: (1) a blood sample, at least one 6 ml blood sample was drawn using a lavender-top Vacutainer tube containing EDTA; (2) a saliva sample collected using Oracollect®•DNA OCR-100 swab (DNA Genotek), designed for non-invasive collection of high-quality DNA from oral cells. The swab and blood tube were both labeled with the participant’s identification number.

### Immunomagnetic PBMCs isolation

Peripheral blood mononuclear cells (PBMCs) were isolated on-site from freshly collected blood samples (6 mL per participant in EDTA vacutainer tubes) within a maximum 4 h post blood draw using the EasySep™ Direct Human PBMC Isolation Kit (STEMCELL Technologies) following the manufacturer’s instructions with modifications to adapt it to field conditions. Two milliliters of whole blood were transferred to a 5 mL (12 × 75 mm) polystyrene tube and mixed with 100 μL of Isolation Cocktail by pipetting 10–13 times (without vortexing or inversion to avoid red blood cell contamination). Sample was incubated for 5 min at room temperature (RT). Then, two milliliters of EasySep buffer were added, mixed gently, and combined with 100 μL of RapidSpheres previously vortexed for 30 s. Tubes (uncapped) were placed in the EasySep magnet for 5 min at RT, and ∼3 mL of enriched suspension was carefully transferred to a new tube. A second depletion step was performed by adding 100 μL of RapidSpheres to the enriched fraction, mixing by pipetting, and repeating the magnetic incubation for 5 min, after which the clear supernatant was collected. A third magnetic incubation of the enriched fraction for 5 min yielded ∼2.5 mL of highly purified PBMC suspension. This negative selection approach, based on antibody-coated magnetic particles that bind and deplete red blood cells and platelets, avoided centrifugation and electrically powered equipment, yielding untouched and functionally intact PBMCs suitable for immediate cryopreservation and downstream applications.

### FDG PBMCs isolation

For the Ficoll method, blood was diluted 1:1 with PBS1x and carefully added to 15 mL conical tube containing 5 mL of Ficoll solution. After a 40 min centrifugation at 500 x g without break, the buffy coat containing the PBMCs were collected into a new 15 mL tube with 5 mL of PBS 1x and centrifuged for 5 min at 500 x g. The cell pellets were washed two more times as previously described. At the end of isolation protocols, the cells from both methods were frozen in inactivated Bovine Fetal Serum (Gibco, Waltham, Massachusetts, USA) with 10% DMSO (LGC, London, United Kingdom).

### Cryopreservation

Following PBMC isolation, cells were cryopreserved in a solution consisting of 10% dimethyl sulfoxide (DMSO) and 90% fetal bovine serum (FBS), prepared by gently adding 0.5 volumes of 30% DMSO in FBS (1.25 mL) to 2.5 mL of purified PBMC suspension, keeping the mixture on ice at all times to minimize DMSO toxicity. The final suspension contained 0.5–10 × 10^6 cells/mL, and three precooled cryovials (CORNING, 2 mL each) were prepared per donor. Cryovials were labeled and arranged in advance. Additional “calibration” vials containing 1 mL of cryomedium were included to ensure all wells of the CoolCell® freezing container (BioCision) were filled, as the device is calibrated for use when full. The CoolCell container was pre-equilibrated on ice and immediately transferred into an insulated box filled with dry ice, allowing controlled-rate cooling (∼1°C/min) without electricity. Samples remained fully submerged in dry ice during transport, with constant replenishment to prevent exposure above –70°C, and precautions were taken to avoid CO_2_ accumulation in enclosed vehicles during handling of >5 kg dry ice. Once equilibrated at –80°C for ≥4 h, cryovials were transported either fully embedded in dry ice, transferred to smaller insulated boxes for shipment, or, when possible, moved into liquid nitrogen tanks for extended storage, where they were kept in vapor phase below –135°C until downstream use.

### DNA Extraction

Saliva samples were collected using Oragene® collection kits (DNA Genotek Inc.), which stabilize and preserve DNA at room temperature for up to six months. DNA was extracted following the manufacturer’s protocol with the prepIT®•L2P Kit (DNA Genotek Inc.), optimized for high-yield recovery from saliva. DNA concentration was measured using a Qubit® Fluorometer (Thermo Fisher Scientific), and purity was evaluated by NanoDrop™ spectrophotometry (Thermo Fisher Scientific), with acceptable A260/A280 ratios ranging from 1.7–2.0. Integrity was assessed by agarose gel electrophoresis to verify the predominance of high–molecular weight DNA. Samples exhibiting strong degradation or abnormal purity ratios were excluded from downstream applications. Only DNA extractions with concentrations >20 ng/μL, total yield >200 ng, and no evidence of contamination (e.g., protein carryover or RNA traces) were advanced to genotyping.

### Microarray Genotyping

Genotyping was carried out with the Infinium® Global Screening Array v2.0 (GSAv2, Illumina), which interrogates ∼650,000 SNPs representative of diverse global populations and enriched for clinically relevant loci. The Infinium workflow comprises DNA amplification, fragmentation, precipitation, hybridization to beadchips, single–base extension, enzymatic staining, and scanning, requiring ∼2–3 days to complete. DNA samples passing initial quality thresholds (≥200 ng input, A260/A280 1.7–2.0, intact high–molecular–weight DNA) were processed according to the manufacturer’s instructions.

### Cell thawing and viability

Cells were recovered from liquid nitrogen storage and kept on dry ice during transport to the bench. Cryovials were immediately thawed in a 37 °C water bath for ∼2 min, or until only small ice fragments remained, taking care not to submerge vials beyond the screw-cap to prevent contamination. Immediately after thawing, the cell suspension was transferred into a pre-labeled 15 mL Falcon tube containing 8 mL of prewarmed (37 °C) thawing buffer (RPMI supplemented with 10% FBS and 0.04% BSA), and centrifuged at 400 g for 5 min at room temperature. The supernatant was carefully aspirated with a pipette, avoiding disruption of the pellet, which typically appeared as an opaque ring at the bottom of the tube. The pellet was washed with 4 mL of washing buffer (1× PBS + 5% FBS + 0.04% BSA), centrifuged again under the same conditions, and the supernatant was discarded. Cells were then slowly resuspended in 0.4 mL of washing buffer and transferred into sterile pre-labeled Eppendorf tubes, yielding a suspension close to the required concentration for downstream single cell applications. Cell concentration and viability were assessed by Trypan Blue exclusion (0.4%), using either manual microscopy-based counting or automated counters, with a minimum viability threshold of 70% required and ∼90% considered optimal. When necessary, concentrations were adjusted to fall within the optimal range of 800–1,200 cells/μL, aiming for 1,200 cells/μL and a final recovery of 20,000 PBMCs per sample pool (5,000 cells per individual). It was preferable to achieve a high concentration at the resuspension step and dilute with washing buffer as needed, rather than perform additional centrifugation steps that could compromise viability. Once the target concentration was achieved, cells were passed through a 40 μm strainer to eliminate aggregates and maintained on ice until single cell encapsulation.

### Cellular Multiplexing, library construction and sequencing

For single cell RNA sequencing, cells from different participants were pooled together based on equal cell ratios. To maximize genetic diversity and minimize processing costs, four individuals were selected for each multiplexed library construction. Cryovials from the selected individuals were thawed, and cells were washed before a viability assessment was performed using a microscope-based cell count. The Chromium Single Cell 3’ Library & Gel Bead Kit (10x Genomics) was used to generate single cell libraries. Each participant contributed approximately 5,000 cells per multiplex, targeting a recovery of around 20,000 cells per well on the 10x Genomics Chip M. After generating gel bead-in-emulsion (GEM) droplets, the reverse transcription and cDNA amplification was performed within the droplets, incorporating unique molecular identifiers (UMIs) and cell barcodes. Library preparation followed the standard 10x Genomics protocol, including fragmentation, end repair, A-tailing, adaptor ligation, and PCR amplification. Quality control of the libraries was performed using Qubit® Fluorometry for quantification and Agilent 2100 Bioanalyzer or TapeStation 4150 capillary electrophoresis for size distribution analysis. Three processing HUBs were established across Latin America. The samples used in this study were processed in Mexico, Chile, and Brazil. Libraries were sequenced using the NovaSeqX Plus platform and the NovaSeq 6000 platform using S4 flowcells and Illumina-recommended protocols.

### VCF generation

DNA sequencing data (from microarrays) was used to perform variant calling using GenomeStudio (v2.0). Variants with CallFreq <95% were removed, and copy number variants (CNVs) annotated as [D/I] or [I/D] were also excluded. After recalculating the sample statistics, only donors with a CallRate greater than 95% were retained. We removed all individuals with >5% missing genotype data and all genotypes with >5% missing individuals. We limited the analysis to biallelic SNP and duplicated SNPs were removed. Subsequently, BIM files were generated with (*29*). The resulting VCF contains merged genotypes from all donors participating in LatinCells (∼650,000 SNPs).

### Mapping and Quality control

Raw scRNA-seq reads were processed using Cell Ranger (v8.3.0, 10x Genomics) with default parameters against the GRCh38-2024-A human reference genome (Ensembl), resulting in the generation of gene-by-cell expression matrices. Mapping quality was assessed through standard Cell Ranger summary metrics, including the fraction of reads confidently mapped to the transcriptome, sequencing saturation, mean reads per cell, and mean genes detected per cell. BAM files were generated for each multiplexed pool for subsequent demultiplexing and downstream analyses.

### Genotype-based Demultiplexing

Demultiplexing of single cell data at the donor level was performed with Demuxlet (v1.0), which leverages genotype information from matched donors to assign each cell to its most likely individual of origin. For this, we filtered our genotype freeze using donor identifiers to generate a pool-specific VCF file for each multiplexed library, ensuring direct compatibility with the scRNA-seq data. All runs were conducted using default parameters as recommended by the developers. To increase confidence and efficiency in genotype-based demultiplexing, we employed Freemuxlet (Popscle suite; https://github.com/statgen/popscle), which performs demultiplexing without requiring matched donor genotypes. Instead, Freemuxlet infers donor identities de novo by clustering cells based on SNP information. For this purpose, we used a reference VCF file derived from the 1000 Genomes Project (29), filtered to include only exonic variants with minor allele frequency (MAF) >5%. This file can be found on the popscle page. The expected number of donors per pool (3-4) was specified, and all analyses were run with default parameters. Finally, we integrated results from both Demuxlet and Freemuxlet to generate consensus donor assignments, maximizing robustness in donor identification across multiplexed pools.

### scRNA-seq data analysis

We excluded low-quality cells using the following thresholds: <15% of mitochondrial RNA percentage, <500 genes per cell, and 800-20,000 UMIs. Gene expression was normalized and corrected using a 10,000 factor. 2,000 variable genes were calculated and scaled, and PCA matrix was inferred. Clusters were generated using the top 25-50 PC dimensions (according to the *ElbowPlot* function on each dataset) and visualized on a UMAP. We finally merged and integrated datasets from both protocols to eliminate batch effects using the CCA algorithm from Seurat.

### Collection of scRNA-seq data generated with FDG method

Raw scRNA-seq data was obtained from Gene Omnibus Expression (Table 1) and was selected based on the following criteria: 1) FDG was the PBMC isolation method, 2) only healthy donors were considered, 3) each donor must have at least 5,000 raw cells, and 4) libraries were constructed using 10X Genomics RNA library kit.

### Analyses of cell quality, relative cell type proportion correlations, and stress genes

To compare the performance of both FDG and SiteCELL protocols, we utilized metrics such as 1) cell quality and changes in cell expression profile, 2) cellular proportion variation and cell contamination, and 3) cellular diversity captured. To assess cell quality we evaluated and calculated percentage of cell viability, UMIs and genes per cell, as well as mitochondrial percentage. Cellular proportions were calculated for both subtypes and major group level in all samples after cell type annotation using a PBMC reference in the Azimuth R library (v0.5.0) (*30*). We annotated cells using the second resolution which identifies 30 different cell types, including other blood cells such as red blood cells and platelets and calculate the relative proportion by dividing the number of cells in each subtype by the total number of cells and multiplying by 100. To evaluate changes in gene expression profiles of matched samples, we integrated all datasets included in the comparison and generated a pseudobulk by summing together the gene counts of all the cells grouped by dataset and major cell type using *AggregateExpression* and then comparing the normalized expression values. To evaluate cell stress, we employed a set of genes previously reported to change due to cellular stress (Massoni-Badossa et al) (*SRGN, SAT1, RBM3, HEAT1, FTH1, EIF1, CXCR4, CIRBP*) and average their expression values per donor and per batch. We replicated this analysis for the experiment assessing performance of SiteCELL in rural settings vs. available FDG experiments using same aggregation strategy but including and expanded set of genes reported to vary due to time delay in processing (Savage et al) (*SRGN, SERF2, SAT1, PFN1, NEAT1, MYL12A, KLF6, JUND, JUNB, GIMAP7, FOS, DUSP1, CFL1* and *CD52*). Lastly, to evaluate rare cell subtype capture efficiency we created a subset of all T cells and then re-clustered and annotated those cells. We re-ran *FindCluster, FindNeighbors* and *RunUMAP* on the subsetted object and calculated cell subtype relative proportions as before. Finally, to calculate the expression level correlation of cell identity genes, we averaged the normalized expression value of each gene across major cell types per donor (from pseudobulked counts) and calculated person correlation for the top 10 unique marker genes per cell subtype (219 in total) across donors from matched experiments.

## Data and code availability

Ficoll Single cell RNA-seq data are publicly available from GEO with the following accession numbers:

- Ficoll 1: GSE226896
- Ficoll 2: GSE149689
- Ficoll 3: GSE161918
- Ficoll 4: GSE213516
- Ficoll 5: GSE168732

- All original code has been deposited at Zenodo at [DOI] and is publicly available as of the date of publication.
- Any additional information required to reanalyze the data reported in this paper is available from the lead contact upon request.

