## Supplementary material for "SiteCELL enables *on-site* PBMCs purification and cryopreservation for immune single cell profiling of diverse ancestries": SiteCELL bench-ready manual

### On-site PBMCs isolation and cryopreservation

#### Principle

PBMC isolation protocols involve the separation of lymphocytes from whole blood samples. Since whole blood is composed primarily of plasma, platelets and red blood cells, PBMCs are extremely difficult to study together with other cell types. Therefore, isolation must be performed before any downstream processing or analyses.

The most common method for PBMCs isolation is density-gradient centrifugation, which relies on the different size and density of PBMCs to separate them from other blood cells. Although this technique is cost-effective and relatively efficient, it´s time consuming and requires specialized laboratory equipment. Therefore, it is not suitable for isolating PMBCs in remote locations without technical equipment and electricity.

#### On-site isolation protocol

This protocol is intended for isolating and cryopreserving PBMCs on-site, with the objective of characterizing their transcriptional state using single-cell RNAseq technology. It has been specially designed for scenarios in which blood samples are collected in remote locations with no access to electricity and far from processing facilities.

Routine protocols for collecting blood with the objective of downstream PBMCs processing recommend that cells should be isolated from blood and cryopreserved within a window of 24 hours from the time of collection. After that time frame, cells start to dramatically lose viability. Therefore, there is currently no protocol available that allows blood collection and PBMCs isolation from remote communities where access is difficult and sample transportation to the closest processing facility could take several days.

#### Key technical aspects

Isolation and purification of PBMCs on-site prevents transcriptional changes and viability issues due to sample transport. Damaged or dead cells upon thawing will interfere with single-cell RNAseq protocols, therefore preserving viability is paramount. Importantly, cryopreservation protocols do not work on whole blood samples, hence PBMCs have to be isolated before cryopreservation can be performed.

***Isolation.*** PBMCs will be purified with an immunomagnetic kit that uses antibodies to select and separate unwanted cells from the sample (negative selection). This will leave only lymphocytes for downstream processing. Antibodies are conjugated to the surface of magnetic beads and targeted to bind red blood cells and platelets. These are then precipitated using a magnet. Therefore, no centrifugation is needed. The protocol, which is described in annex 1 basically consists of a series of washing steps.

***Cryopreservation.*** Successful freezing of PBMCs depends on the addition of a cryoprotectant reagent like DMSO to the cell mix. This prevents the formation of crystals during the freezing process. In addition, to preserve cell viability it is very important that temperature during freezing decreases at a rate of 1 $C^o$ per minute until reaching -80 $C^o$. A cheap and convenient way to achieve this is by using CoolCell containers, which have been specially designed with internal microcirculation that allows gradual freezing. When done in the lab, cryovials containing freezing medium and PBMCs are placed on coolcell containers and moved to a -80 freezer. However, in the field this step can be replaced by placing the containers in an ice box filled with dry ice, that has an approximated temperature of -79 $C^o$.

***Long-term storage.*** After 10 days at -80 $C^o$, cryopreserved cells can start developing crystals which greatly impact viability; hence, they should be transferred to a liquid nitrogen container for long term storage. At a temperature of -180 $C^o$ cells can be preserved for years without loss of viability.

#### Stage 1: Logistics.

Although protocols have been simplified to allow processing on remote locations, the number of materials and the degree of coordination needed is comparable to mounting a small, mobile laboratory on-site. As a result, the success of sample collection depends on careful planning before traveling to the collection site. A few parameters have to be defined before every collection trip:

1. The number of samples that will be collected. In other words, how many donors do you expect.
2. The number of staff available for the sampling trip.
3. Expected time required to get into the community and out to a processing hub or shipping station.
4. Mode of transportation and number of vehicles.
5. Expected weather during sample transport.
6. Services available near the collection site: will you have access to regular ice and running water?

##### *1.1 Material*

Having a good idea of the number of samples you intend to collect is critical because this will determine the material and equipment you need to organize and transport. Once on-site, this will be impossible to adjust. Annex 2 provides a list of indispensable materials that need to be acquired and transported based on the number of samples/donors.

Most of the reagents, consumables and equipment has to be acquired and organized several weeks before the sampling trip, but some items like dry ice, and cryopreservation media are prepared on the previous day. See the “details” section on annex 2. It’s important to comply with the technical specifications of all materials, including brand, quantity, and size. If a material or equipment does not work on site, it will be impossible to replace it on time, and the trip will have to be canceled at that point. Therefore, only materials tested in pilot studies are recommended.

***Dry Ice.*** Getting and transporting enough dry ice for sampling trips is a logistical challenge and should be carefully planned. Dry ice will be needed for PBMCs cryopreservation on-site. Once frozen, the cells have to be kept inside the coolcell containers and covered by dry ice at all times during transport. If cells get thawed before they are processed for single cell library prep they have to be discarded. Please consider the following information to calculate the amount of dry ice needed and transportation requirements:

a) Dry ice will sublimate at a rate of 4.5-5 kgs every 24 hours. Therefore 20 kgs will last for 4 days until completely evaporated. However, if exposed to warm conditions this rate could increase substantially. In addition, consider that the point of complete evaporation should never be reached. You should always calculate in excess.

b) In a pilot conducted in México where the average temperature at that location was 20-25 Co, 40kgs of dry ice were necessary to cryopreserve cells and to keep them completely covered for three days. This is an average of 13 kgs/day. For longer periods of time more dry ice has to be transported or a resupply shipment to the sampling point must be arranged.

c) The size of the Ice-box (insulating container) where the dry ice will be stored and transported will also impact the amount of dry ice needed. We recommend using a container having the following approximate size: 21.5 inch wide, 18.5 inch high, and 17 inch deep. This will allow to accommodate up to four coolcell containers, with space for 30 tubes each. The amount of dry ice specified above is based on an ice-box with this volume and will ensure coolcell containers will be always covered with dry ice.

d) We strongly advice to transport a back-up dry-ice load in case the dry ice evaporates faster or in the event of a spill.

**1.2 Staff**

During the day of sample collection, we recommend organizing activities in 4 stations, operated by a total of 6 persons (minimum):

- *Consent and survey.* We recommend two people being in charge of fill out questionnaires and explaining consent forms to participants. This can be time consuming and a delay can impact the rate of PBMCs processing downstream.

- *Biometrics.* One person in charge of taking participant´s biometrics.

- *Blood collection*. One person should be focused solely on taking blood samples.

- *PBMCs isolation*. No less than two persons should be handling PBMCs isolation protocol. Samples have to be processed and cryopreserved in batches and there is considerable complexity and logistics in the protocol.

* If less than 15 samples will be collected, the staff could be reduced.

**1.3 Transportation time**

Considering transportation time is important because it directly impacts the amount of dry ice that needs to be arranged. If reaching the community takes more than two days (4 days in total to get back) a bigger transport should be arranged to accommodate staff and several dry ice containers. * Please calculate at least 13 kgs of dry ice per day, plus backup.

##### 1.4 Services available

Regular ice will be needed for the cryopreservation protocol. Therefore, if access to a convenience store is restricted take several ice packs, as small as possible (see annex 2). The ice packs can be kept frozen if stored in the dry ice container until use.

Because you will be handling potentially hazardous biological material, disinfection will be important. Therefore, if running water is not available, sufficient bleach and disinfecting wipes or similar have to be considered.

***Important note:***

Material and logistics checklists are provided. Ensure you fulfill all requirements enlisted before starting your sampling trip.

#### Stage 2: PBMCs isolation and cryopreservation

A considerable number of materials and reagents are needed during the isolation and cryopreservation process. All materials should be organized before starting and located comfortably at hand before starting the protocol. It is recommended to use at least two medium-sized portable tables and chairs to set up a working station.

##### 2.1 sample processing

Because Coolcell containers can accommodate 30 cryovials and cannot be reopened once placed on dry ice, blood samples have to processed in batches. Therefore, if 3 PBMCs cryovials are obtained per donor, ideally you should process a 10-sample batch in one go to completely fill the container. Less samples per batch can be processed, but Coolcell space will be wasted. We advise waiting until 10 blood samples have been collected to start the isolation protocol. Blood samples can be left at room temperature for up to 4 hours without considerable transcriptome changes or loose of viability.

***BIOHAZARD WARNING.*** Blood samples pose a potential biohazard, therefore caution should be taken during blood collection and PBMCs isolation. Direct contact with blood or plasma should be avoided and any spills should be properly disinfected. Do not attempt to isolate PBMCs if you are not trained in handling blood. You should wear the following protective gear as a minimum when processing samples:

1. Long lab coat.
2. Nitrile gloves.
3. N95 face mask or equivalent.
4. Protective lenses that cover most of your cheeks.

Proceed to PBMCs isolation protocol annex 1 (below) when all previous considerations have been covered.

##### 2.2 Cryopreservation

The process of cryopreservation consists of three basic steps: **a)** mixing the isolated PBMCs with the appropriate concentration of cryoprotectant (DMSO) and fetal bovine serum (FBS), **b)** transferring cryovials to coolcell containers, and **c)** organize coolcell containers inside an insulating box filled with dry ice.

Successful cryopreservation depends on the speed at which samples are prepared and transferred to freezing conditions. Because DMSO is toxic to cells at room temperature, once it has been added to the cell mix is very important to keep cells on ice and minimize as much as possible the time it takes to prepare the vials and transfer them to coolcell containers and dry ice.

We recommend labeling, organizing, and precooling all cryovials (3 per donor) before adding DMSO to isolated PBMCs. We also recommend preparing several “calibration” cryovials beforehand. These are filled with 1 ml cryopreservation media (at least 10 vials). They will be needed to fill empty spaces in the coolcell container when small blood sample batches are prepared. Cryopreservation medium should be prepared in the lab before traveling and transported frozen to the site inside the dry ice box.

Once everything is set, please follow the cryopreservation protocol below (annex 3)

#### Stage 3: Transportation and thawing

Once cells have been cryopreserved, they cannot be thawed during transport. Cells that are exposed to temperatures above -70 C$^{o}$ will have to be discarded. Therefore, it is essential to always keep coolcell containers completely covered with dry-ice and transport them as soon as possible to the closest shipping point or processing hub.

##### 3.1 Dry-ice transportation

Insulation boxes containing dry ice and samples will have to be transported by land, at least for a segment of the journey. When handling considerable amounts of dry-ice in a small space, care has to be taken to provide sufficient ventilation, as dry ice will produce carbon dioxide gas when it sublimates.

For example, 40 kgs of dry ice transported inside a car will quickly raise the CO2 concentration to dangerous levels if windows are closed. Since CO2 is odorless, a driver can faint before realizing something is wrong. Therefore, the following recommendations have to followed when transporting samples in dry ice:

1. Always use and store in a ventilated area, especially if employing more than 5kgs.
2. If possible, keep it in a cool place and protected from direct sun.

3. When transported in a vehicle, keep windows open at all times.
4. Don’t store dry ice in an airtight container.
5. Wear adequate protective gear to handle dry ice.

To avoid sample temperature from rising, constantly verify that coolcell containers are completely covered with dry ice. If you start to see the top of the container, immediately refill with backup dry ice. **Once cells are completely frozen and equilibrated to -80 (allow at least 4 hours),** you have a few options that can potentially extend transportation time:

a) If needed, cryovials can be quickly removed from the coolcell container and placed directed in dry ice. Less dry ice is needed to keep the vials completely covered compared to the bulky coolcell containers. Empty coolcell containers can be removed from dry ice to create more space.

   *Note: Only do this if really necessary, to avoid exposing cells to temperature changes.*

b) Cells can be quickly removed from coolcell containers and placed on a liquid nitrogen tank for transportation. Some nitrogen tanks are very well insulated and last for several days up to a week. However, once in liquid nitrogen cells cannot go back to dry ice, they have to be directly transferred to stable cryo-tank for long term storage.

c) If you are close to a shipping point, cells could be quickly removed from the coolcell container and placed directly on dry ice in a smaller insulting box appropriate for shipping. Samples could be shipped by air directly to a processing hub.

**3.2 Thawing**

*This section is only relevant if you are a processing hub.*

For processing samples with chromium 10x for single-cell RNA profiling cells will have to be recovered from liquid nitrogen storage and thawed. The protocol consists of two steps: rapid thawing at 37 Co and a couple of washing and centrifugating steps.

Note: it is important to remember that cells are still in DMSO containing media, therefore once thawed cells have to centrifugated and washed as soon as possible.

Follow the complete thawing protocol below (annex 4).

**This bench-ready protocol version has been prepared as a supplemental material from the publication “SiteCELL enables on-site PBMCs purification and cryopreservation for immune single cell profiling of diverse ancestries”.*

### Annex1: isolation of PBMCs with Immunomagnetic kit

Purification of PBMCS will be performed by immunomagnetic negative selection using the *EasySep Direct Human PBMC Isolation Kit.* This kit uses magnetic beads attached to antibodies that bind and precipitate unwanted cells (red blood cells and platelets), eliminating the need for centrifugation. Steps are as follow:

*Notes:*

- *It is crucial that 6 milliliters of blood are collected in EDTA vacutainer tubes. This will ensure appropriate EDTA concentration for downstream purification protocol.*
- *Always pipette mix by pushing the pipette push button 10-13 times.*

1. Transfer 2ml of recently collected blood (no more than 4 hours since collection) to a 5ml (12 x 75mm) polystyrene tube.

2. Add 100 µl of Isolation cocktail to the sample (don't vortex cocktail).

3. Mix with pipette approximately 10 times. ***IMPORTANT:*** don't invert the tube as this will cause red blood cell contamination!

4. incubate for 5 min.

5. Add 2ml of EasySep buffer. Mix by gently pipetting up and down 2-3 times.

6. Vortex RapidSpheres for 30 seconds.

7. Add 100 µl of RapidSpheres to sample and mix by pipetting. Immediately proceed to next step.

8. Place the tube without lid into the magnet and incubate for 5 min at RT.

9. Carefully pipette ≈ 3 ml of the enriched cell suspension into a new 5 ml tube. ***IMPORTANT*:** Take care not to touch the tube opening with the pipette tip as it may contain red blood cell contamination from step 1.

10. Add 100 µl of RapidSpheres into the new tube containing the enriched cells and mix by pipetting.

11. Place the tube from step 9 into the magnet and incubate for 5 min for a second separation.

12. Carefully pipette the enriched cell suspension into a new 5 ml tube. Collect only the clear fraction.

13. Place the tube from step 11 into the magnet and incubate for 5 min at RT for a third separation.

14. Carefully pipette 2.5 ml of the purified cell suspension into a new tube. Isolated cells are ready for cryopreservation protocol.

### Annex 2: Material Checklist

* Calculated for procesing approx. 20 blood samples

#### 1. Blood collection

| Reagent/consumable | Quantity | Transport @ | Check |
|---|---|---|---|
| Vacuteiner EDTA, 6ml | 30 | RT | |
| Vacuteiner needles, 0.8 x 32 mm | 30 | RT | |
| cotton swabs | 2 packs | RT | |
| Alcohol | 2 small bottles | RT | |
| Small gloves | 1 pack | RT | |
| Medium gloves | 1 pack | RT | |
| Big gloves | 1 pack | RT | |
| Tourniquet | 1 | RT | |
| Tray for 6 ml tubes | 4 | RT | |
| Timer | 2 | RT | |
| Sample labels | 20 | RT | |
| Solid waste container | 1 | RT | |
| Liquid waste container | 1 | RT | |
| Vacuteiner adaptor | 3 | RT | |
| Wipes | 3 packs | RT | |
| Bleach | 1 bottle | RT | |
| Protection glasses | 3 | RT | |
| Lab coat | 3 | RT | |
| Big container (4 gallons) | 1 | RT | |
| Biohazard plastic bags | 1 pack | RT | |
| Tape | 1 | RT | |
| Sharpies | 4 | RT | |

#### 2. Isolation of PBMCs

| Reagent/consumable | Quantity | Transport @ | Check |
|---|---|---|---|
| 5 mL (12 x 75 mm) polystyrene tubes | 80 | RT | |
| Isolation cocktail (stemcell) | 1 tube | 4 oC | |
| EasySep Buffer | 1 bottle, at least 30ml | 4 oC | |
| RapidSpheres (stemcell) | 1 tube | 4 oC | |
| Magnet | 1 easy eights | RT | |
| MicroPippettes (1ml, 0.2 ml, 0.02ml) | 2 each | RT | |
| Pippete tips (1ml, 0.2 ml, 0.02ml) | 2 box each | RT | |
| 10ml Serological pippetes | 1 bag | RT | |
| Pippete gun | 1 | RT | |
| 15 ml Falcon tubes | 40 | RT | |
| 1.5 ml microcentrofuge tubes | 30 | RT | |
| Ice | 10-20 kgs | | |
| Ice box | 2 | RT | |
| 2 ml tube rack | 3 | RT | |

| | | |
|---|---|---|
| Batery Vortex | 1 | RT |

#### 2. Cryopreservation of PBMCs

| Reagent/consumable | Quantity | Transport @ | Check |
|---|---|---|---|
| Fetal Bovine Serum (FBS) | 60 ml | -20 | |
| DMSO | 30 ml | -20 | |
| CoolCell container | 2 | RT | |
| 2ml Flat-bottom tubes (Cryopreservation) | 60 | RT | |
| Control vials (15% DMSO, 45% FBS) 1ml | 30 | -20 | |
| Freezing medium (30% DMSO in FBS) | 120 ml | Prepare fresh | |
| Dry Ice | 40 kgs | | |
| Falcon tubes 50 ml | 20 | RT | |

### Annex 3: cryopreservation in 10% DMSO with 90% FBS

**Principle**

Peripheral blood mononuclear cells (PBMC) can be frozen and stored in a cryoprotective media containing 10% dimethyl sulfoxide (DMSO) and fetal bovine serum (FBS), then thawed rapidly. Cells frozen and thawed in this manner should have an acceptable yield and viability post-storage.

**Before You Begin:**

- Ensure all media is cold prior to starting this protocol.
- Prepare 30% DMSO in FBS. Keep on ice. Note: Do not put 100% DMSO on ice or it will form crystals.
- Label cryogenic vials (3 per donor).
- Ensure PBMCs are in a single-cell suspension.

1. Add gently 0.5 volumes of 30% DMSO in FBS to the cell suspension (that is 1.25ml of cryo medium to 2.5ml of cell suspension). The final cell suspension will be in 10% DMSO and 90% FBS. The final cell concentration will be between 0.5 - 10 x $10^6$ cells/mL.

   ***Note: Do not let cells sit in cryopreservation medium at room temperature. Keep on ice and transfer rapidly.***

2. Transfer 1 mL of cell suspension to each cryovial (pre-cooled) and ***keep on ice until freezing***. This means a minimum of 3 cryovials per sample/donor.

3. Place cryogenic vials inside the coolcell container. The Coolcell container should be at the same temperature as the samples. Therefore, place the container on ice and allow it to equilibrate before transferring samples.

   *Note: Fill all the container wells. Coolcell is “calibrated” for use when full. If you don´t have enough samples, use “mock” cryogenic tubes filled with DMSO and FBS.*

4. Immediately transfer the filled coolcell container to the the ice box and completely cover it with dry ice. It is best to slip the container inside a tightly adjusted cardboard box and then cover with dry ice. This will prevent direct contact with dry ice, which can eventually damage the container.

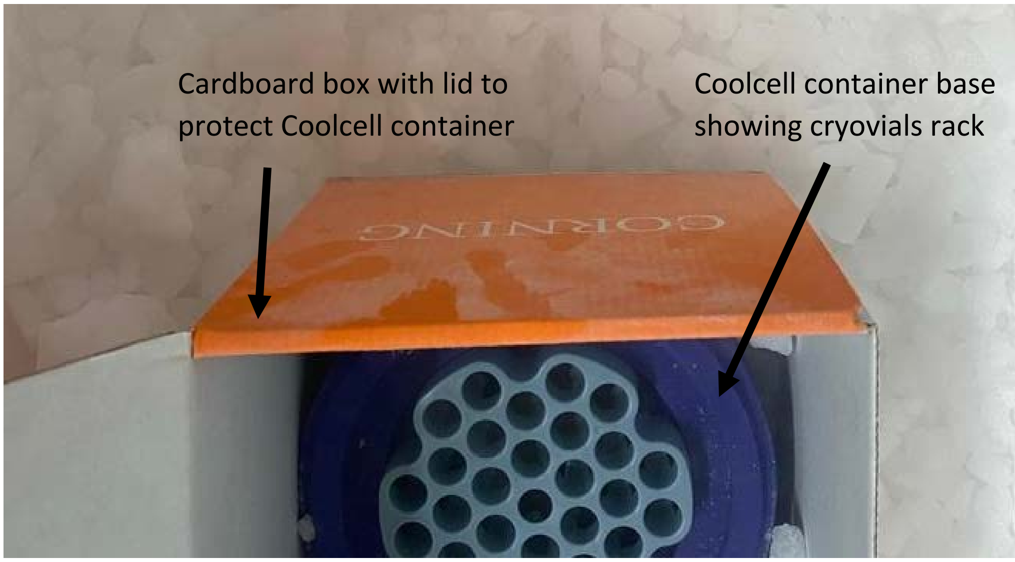

*Image 1. Coolcell container inside cardboard box before adding cryovials with cells.*

**Note:** Once the coolcell container is closed and has been covered by dry ice it cannot be re-opened to shift or accommodate more samples. This can have a big effect on cell viability. Only re-open the container when transferring cells to liquid nitrogen for long term storage.

#### Liquid nitrogen storage

For long-term storage (more than 10 days), transfer the vials with frozen PBMCs to vapor phase liquid nitrogen (below -135°C). Avoid exposure to room temperature by placing coolcell container and vials on dry ice during transfer to liquid nitrogen.

### Annex 4: Cell Thawing & GEM Generation.

#### Before You Begin:

- Prepare thawing buffer (RPMI + 10% FBS + 0.04% BSA). Sterilize by filtration and add **8 mL to 50ml sterile falcon tubes**. Prepare as many as the number of samples that will be processed.
- Prepare washing buffer (1X PBS + 5% FBS + 0.04% BSA). Sterilize by filtration.
- Plan which samples will be processed, locate them, and label the falcon tubes previously prepared.

  *Buffers can be prepared one day before and stored at 4°C.*
- **Prepare water bath at 37 °C**
- Prewarm thawing media to 37 °C
- Book the centrifuge and assemble the rotor.
- Label sterile eppendorf tubes with the sample names (2 per sample)
- Keep washing buffer at 4 °C

*Tip: When dealing with cell suspensions always pipette very slowly and use wide bore pipettes if possible.*

**1.** Retrieve the cells from liquid nitrogen storage and place them directly on **dry ice** while transporting them to the bench.

**2.** Place the vials directly in a water bath or similar at 37 °C for approximately two minutes, or until very little ice remains. If using a water bath, avoid introducing the vials beyond the screw cap level as water may get into the tube.

**3.** After thawing, immediately transfer the cells into a 50ml falcon containing 8 ml of pre/warmed (37 °C) thawing media.

**4.** Centrifuge the tubes at 400 g for 5 minutes (room temperature).

**5.** Remove the supernatant with a pipette taking care not to disturb the cell pellet.

*Note: If using 50 ml falcon tubes, the cell pellet might be difficult to see as it will form a ring shape at the bottom of the tube. Check carefully for the presence of an opaque ring at the bottom and take care not to disturb it when removing the media.*

**6.** Add **4 mL of tawing media** to the pellet without disturbing it and repeat steps 4 and 5.

**7.** Slowly resuspend the cells in 0.4 ml of washing buffer and transfer them to an eppendorf tube.

*Note: resuspending in 0.4 ml will result in a cell concentration that will be in close range to the concentration needed as input for a chromium single-cell run.*

**8.** Proceed to evaluate the cell concentration and viability.

💡 At this point a lab mate should retrieve the following GEM reagents either form the -80 °C or from the -20°C and equilibrate them at room temperature.

Single Cell 3' HT v3.1 Gel Beads (--80 °C)
RT Reagent B (--20 °C)
Template Switch Oligo (--20 °C)
Reducing Agent B (--20 °C)

A **cell viability above 70%** is required for single-cell experiments. Viability percentages around 90% are desirable.

**9.** Readjust the cell concentration if necessary. The optimal concentration for *single-cell* profiling is from 800 to 1200 cells/µL. For the PBMCs experiments we are aiming for a concentration of **1200 cells/µL** and a total cell output of 30,000 cells (see page #3 for calculation details).

*Note: it is best to obtain a high cell concentration from step 7 and dilute as needed using washing medium, as opposed to doing an extra centrifugation step to increase concentration.*

**10.** Once the desired concentration is achieved, pass the cells through a **40 µm cell strainer** to remove potential cell clumps.

**11.** Place the cells on ice until processing.

**12.** Immediately proceed to load the 10x genomics chip. Cells lose viability with time.

#### Cell Suspension Volume Calculator

Volume of Cell Suspension Stock per reaction (µl) | Volume of Nuclease-free Water per reaction (µl)

DO NOT add nuclease-free water directly to single cell suspension. Add nuclease-free water to the Master Mix. Refer to step 1.2c.

| Cell Stock Conc. (Cells/µl) | Targeted Cell Recovery | | | | | | | | | | |
|---|---|---|---|---|---|---|---|---|---|---|---|
| | 2000 | 4000 | 6000 | 8000 | 10000 | 12000 | 14000 | 16000 | 18000 | 20000 | 30000 |
| 100 | 31.3 / 55.1 | 63.2 / 23.2 | n/a | n/a | n/a | n/a | n/a | n/a | n/a | n/a | |
| 200 | 15.6 / 70.8 | 31.6 / 54.8 | 47.8 / 38.6 | 64.1 / 22.3 | 80.7 / 5.7 | n/a | n/a | n/a | n/a | n/a | |
| 300 | 10.4 / 76.0 | 21.1 / 65.3 | 31.8 / 54.6 | 42.7 / 43.7 | 53.8 / 32.6 | 64.9 / 21.5 | 76.2 / 10.2 | n/a | n/a | n/a | |
| 400 | 7.8 / 78.6 | 15.8 / 70.6 | 23.9 / 62.5 | 32.1 / 54.3 | 40.3 / 46.1 | 48.7 / 37.7 | 57.2 / 29.2 | 65.8 / 20.6 | 74.4 / 12.0 | 83.2 / 3.2 | |
| 500 | 6.3 / 80.1 | 12.6 / 73.8 | 19.1 / 67.3 | 25.6 / 60.8 | 32.3 / 54.1 | 39.0 / 47.4 | 45.7 / 40.7 | 52.6 / 33.8 | 59.6 / 26.8 | 66.6 / 19.8 | |
| 600 | 5.2 / 81.2 | 10.5 / 75.9 | 15.9 / 70.5 | 21.4 / 65.0 | 26.9 / 59.5 | 32.5 / 53.9 | 38.1 / 48.3 | 43.8 / 42.6 | 49.6 / 36.8 | 55.5 / 30.9 | |
| 700 | 4.5 / 81.9 | 9.0 / 77.4 | 13.6 / 72.8 | 18.3 / 68.1 | 23.0 / 63.4 | 27.8 / 58.6 | 32.7 / 53.7 | 37.6 / 48.8 | 42.5 / 43.9 | 47.6 / 38.8 | |
| 800 | 3.9 / 82.5 | 7.9 / 78.5 | 11.9 / 74.5 | 16.0 / 70.4 | 20.2 / 66.2 | 24.4 / 62.0 | 28.6 / 57.8 | 32.9 / 53.5 | 37.2 / 49.2 | 41.6 / 44.8 | |
| 900 | 3.5 / 82.9 | 7.0 / 79.4 | 10.6 / 75.8 | 14.2 / 72.2 | 17.9 / 68.5 | 21.6 / 64.8 | 25.4 / 61.0 | 29.2 / 57.2 | 33.1 / 53.3 | 37.0 / 49.4 | |
| 1000 | 3.1 / 83.3 | 6.3 / 80.1 | 9.6 / 76.8 | 12.8 / 73.6 | 16.1 / 70.3 | 19.5 / 66.9 | 22.9 / 63.5 | 26.3 / 60.1 | 29.8 / 56.6 | 33.3 / 53.1 | 1000 cells/µl: 49.9 / 36.5 |
| 1100 | 2.8 / 83.6 | 5.7 / 80.7 | 8.7 / 77.7 | 11.7 / 74.7 | 14.7 / 71.7 | 17.7 / 68.7 | 20.8 / 65.6 | 23.9 / 62.5 | 27.1 / 59.3 | 30.3 / 56.1 | |
| 1200 | 2.6 / 83.8 | 5.3 / 81.1 | 8.0 / 78.4 | 10.7 / 75.7 | 13.4 / 73.0 | 16.2 / 70.2 | 19.1 / 67.3 | 21.9 / 64.5 | 24.8 / 61.6 | 27.7 / 58.7 | 1200 cells/µl: 41.5 / 44.9 |
| 1300 | 2.4 / 84.0 | 4.9 / 81.5 | 7.3 / 79.1 | 9.9 / 76.5 | 12.4 / 74.0 | 15.0 / 71.4 | 17.6 / 68.8 | 20.2 / 66.2 | 22.9 / 63.5 | 25.6 / 60.8 | |
| 1400 | 2.2 / 84.2 | 4.5 / 81.9 | 6.8 / 79.6 | 9.2 / 77.2 | 11.5 / 74.9 | 13.9 / 72.5 | 16.3 / 70.1 | 18.8 / 67.6 | 21.3 / 65.1 | 23.8 / 62.6 | |
| 1500 | 2.1 / 84.3 | 4.2 / 82.2 | 6.4 / 80.0 | 8.5 / 77.9 | 10.8 / 75.6 | 13.0 / 73.4 | 15.2 / 71.2 | 17.5 / 68.9 | 19.9 / 66.5 | 22.2 / 64.2 | |
| 1600 | 2.0 / 84.4 | 4.0 / 82.4 | 6.0 / 80.4 | 8.0 / 78.4 | 10.1 / 76.3 | 12.2 / 74.2 | 14.3 / 72.1 | 16.4 / 70.0 | 18.6 / 67.8 | 20.8 / 65.6 | |
| 1700 | 1.8 / 84.6 | 3.7 / 82.7 | 5.6 / 80.8 | 7.5 / 78.9 | 9.5 / 76.9 | 11.5 / 74.9 | 13.5 / 72.9 | 15.5 / 70.9 | 17.5 / 68.9 | 19.6 / 66.8 | |
| 1800 | 1.7 / 84.7 | 3.5 / 82.9 | 5.3 / 81.1 | 7.1 / 79.3 | 9.0 / 77.4 | 10.8 / 75.6 | 12.7 / 73.7 | 14.6 / 71.8 | 16.5 / 69.9 | 18.5 / 67.9 | |
| 1900 | 1.6 / 84.8 | 3.3 / 83.1 | 5.0 / 81.4 | 6.7 / 79.7 | 8.5 / 77.9 | 10.3 / 76.1 | 12.0 / 74.4 | 13.8 / 72.6 | 15.7 / 70.7 | 17.5 / 68.9 | |
| 2000 | 1.6 / 84.8 | 3.2 / 83.2 | 4.8 / 81.6 | 6.4 / 80.0 | 8.1 / 78.3 | 9.7 / 76.7 | 11.4 / 75.0 | 13.2 / 73.2 | 14.9 / 71.5 | 16.6 / 69.8 | |

**Grey boxes:** Exceeds allowable volume

**Blue boxes:** Optimal cell stock conc. for cell recovery target of 2,000-20,000

**Yellow boxes:** Low transfer volume that may result in higher cell load variability

The volume of cell suspension stock / nuclease free water required for a targeted recovery of 30,000 cells were calculated assuming a **60%** recovery rate approx. Importantly, cell concentrations are maintained in the suggested range of either 1000 or 1200 cells/µl. This will help avoiding an increase in the percentage of doublet generation.

### Annex 5: Cell processing checklist

#### 1. Previous to procesing day (prep for thawing)

| Action | Reference | Check |
|---|---|---|
| Determine how many samples will be thawed and from which genotypes. | LatinCells´ PBMCs sample reference list | |
| Book microscopes and centrifuges to be used. | | |
| Make sure to have enough cell countig chambers. | | |
| Make sure to have enough 1ml pipettes. | | |
| Make dry ice arrangements to have enough for transporting samples from liquid nitrogen to lab. | | |
| Check tryptan blue is ready to be used (diluted to working concetration). | | |
| Have enough hand counters ready for counting cells over the microscope. | | |
| Prepare and filter 50% glycerol if needed (22 micrometer filters). | | |

#### 2. On the same day

| Action | Reference | Check |
|---|---|---|
| Prepare and filter thawing buffer (22 micrometer filters). | Annex 4, LatinCells protocol | |
| Prepare and filter washing buffer (22 micrometer filters). | Annex 4, LatinCells protocol | |
| Obtain all 10X genomics reagents and materials. | 10x single cell 3´ HT reagent kits v3.1 (dual index) | |
| Equilibrate reagents to appropiate temperature according to manual before start. | 10x single cell 3´ HT reagent kits v3.1 (dual index) | |
| Make sure to have microscopes and centrifuges ready. | | |
| Equilibrate water bath to 37 oC. | | |
| Clean working space thoroughly. | | |
| Have cell concentration calculation formulas at hand. | | |

#### 3. Define main roles

| Role | Notes | Check |
|---|---|---|
| Who is centrifuging and washing cells? | More than one person | |
| Who will prepare cell counting alicuots (mixing cells with tryptan blue)? | One person | |
| Who will prepare cell multiplexing mix? | One person | |
| Who prepares enzyme master mix and loads the chip? | Just one person | |
